# Environmental drivers of plankton protist communities along latitudinal and vertical gradients in the oldest and deepest freshwater lake

**DOI:** 10.1101/2020.09.26.308536

**Authors:** Gwendoline M. David, David Moreira, Guillaume Reboul, Nataliia V. Annenkova, Luis J. Galindo, Paola Bertolino, Ana I. López-Archilla, Ludwig Jardillier, Purificación López-García

**Affiliations:** Ecologie Systématique Evolution, Centre National de la Recherche Scientifique - CNRS, Université Paris-Saclay, AgroParisTech, Orsay, France; Limnological Institute, Siberian Branch of the Russian Academy of Sciences, Irkutsk, Russia; Departamento de Ecología, Universidad Autónoma de Madrid, Madrid, Spain

**Keywords:** Lake Baikal, protist, 18S rRNA gene metabarcoding, marine-freshwater transition, light stratification, network analysis

## Abstract

Identifying which abiotic and biotic factors determine microbial community assembly is crucial to understand ecological processes and predict how communities will respond to environmental change. While global surveys aim at addressing this question in the world’s oceans, equivalent studies in large freshwater systems are virtually lacking. Being the oldest, deepest and most voluminous freshwater lake on Earth, Lake Baikal offers a unique opportunity to test the effect of horizontal versus vertical gradients in community structure. Here, we characterized the structure of planktonic microbial eukaryotic communities (0.2-30 µm cell size) along a North-South latitudinal gradient (∼600 km) from samples collected in coastal and pelagic waters and from surface to the deepest zones (5-1400 m) using an 18S rRNA gene metabarcoding approach. Our results show complex and diverse protist communities dominated by alveolates (ciliates and dinoflagellates), ochrophytes and holomycotan lineages, with cryptophytes, haptophytes, katablepharids and telonemids in moderate abundance and many low-frequency lineages, including several typical marine members, such as diplonemids, syndinians and radiolarians. Depth had a strong significant effect on protist community stratification. By contrast, the effect of the latitudinal gradient was marginal and no significant difference was observed between coastal and surface open water communities. Co-occurrence network analyses showed that epipelagic communities are much more interconnected than meso- and bathypelagic communities and suggest specific biotic interactions between autotrophic, heterotrophic and parasitic lineages that influence protist community structure. Since climate change is rapidly affecting Siberia and Lake Baikal, our comprehensive protist survey constitutes a useful reference to monitor ongoing community shifts.

**Originality and Significance Statement:** Lake Baikal is the oldest, deepest and most voluminous freshwater lake on Earth, offering a unique opportunity to test the effects of horizontal versus vertical gradients on microbial community structure. Using a metabarcoding approach, we studied planktonic microbial eukaryotes from Baikal water columns (5 up to 1,400 m depth) across a North-South latitudinal gradient (∼600 km), including coastal and pelagic areas. Our results show that depth has a strong effect on protist community assemblage, but not latitude (minor effect) or coastal vs. open water sites (no effect). Co-occurrence analyses also point to specific biotic interactions as drivers of community structure. This comprehensive survey constitutes a useful reference for monitoring active climate change effects in this ancient lake.

## Introduction

Of all ecosystems, freshwater reservoirs are the most dynamic and concentrate a high biodiversity (Rolls et al., 2018). Freshwater ecosystems are particularly vulnerable to climate change owing to a higher exposure and sensitivity to increasing temperature and other altered conditions, limited dispersal across these fragmented habitats and little-known, but likely modest, resilience potential (Woodward et al., 2010; Markovic et al., 2017). Since microorganisms are crucial in biogeochemical cycles, the impact of climate change will strongly depend on how they will respond to environmental challenge (Cavicchioli et al., 2019). Permafrost-covered areas in the Arctic region (Schuur et al., 2015) and forest-steppe ecotones in Siberia are among the most heavily impacted regions by global warming (Mackay et al., 2017). This includes Lake Baikal, in southern Siberia, which is the oldest (ca. 30 Myr), deepest, and most capacious freshwater lake on Earth (Müller et al., 2001). Lake Baikal is rapidly changing, as can be told from trends in hydrological and hydrochemical processes (Moore et al., 2009; Shimaraev and Domysheva, 2012). The lake sediments represent a continuous record of past climate for over 12 million years (Kashiwaya et al., 2001; Prokopenko et al., 2002) such that Lake Baikal is a unique model to understand and predict microbial community change and how this is linked to carbon cycling and hydrological processes.

A mandatory prerequisite for such a task is to have comprehensive information about the existing microbial community structure. However, if the broad biodiversity of Lake Baikal metazoans, including many endemisms (1455 out of 2595 species described), has been amply documented in the past two centuries, that of microbial life is highly fragmentary. One of the reasons relates to the large dimensions of the lake, which is around 640 km long, attains a depth of ca. 1650 m and contains around 20% of the Earth’s unfrozen freshwater (Sherstyankin et al., 2006; UNDP-GEF, 2015). This, together with its geographical location and its association to a rifting zone make Lake Baikal unique and listed as UNESCO World Heritage Site (UNDP-GEF, 2015). The lake is divided in three major basins (Northern, Central, Southern) by, respectively, the Academician Ridge and the Selenga river delta (Mats and Perepelova, 2011). Its surface freezes in winter for several months, favoring coastal downwelling and deep-water oxygenation (Schmid et al., 2008; Moore et al., 2009). As a result, Lake Baikal ultra-oligotrophic waters are globally cold (∼4°C) and oxygen-rich down to the bottom (Schmid et al., 2008; Moore et al., 2009; Shimaraev and Domysheva, 2012; Troitskaya et al., 2015). Baikal also uniquely hosts methane hydrates, which are stabilized by the low temperatures and high pressures (De Batist et al., 2002; Granin et al., 2019). All these features make Lake Baikal akin a freshwater sea.

Microbial diversity in Lake Baikal plankton was first studied by classical observation and cultural approaches (Maksimova and Maksimov, 1972; Maksimov et al., 2002; Bel’kova et al., 2003) before molecular tools started to be applied at the beginning of the century (Glöckner et al., 2000) and expanded more recently with the generalization of high-throughput sequencing. Several 16S rRNA gene-based metabarcoding studies have targeted pelagic bacteria diversity (Kurilkina et al., 2016; Belikov et al., 2019; Zakharenko et al., 2019; Wilburn et al., 2020) and, more recently, metagenomic analyses have been used to characterize planktonic prokaryotic communities from sub-ice (Cabello-Yeves et al., 2018) and deep waters (Cabello-Yeves et al., 2020), virus-bacteria assemblages in coastal waters (Butina et al., 2019) or viruses from the pelagic zone (Potapov et al., 2019). Microbial eukaryotes have only been partially studied by 18S rRNA gene metabarcoding. Several of these studies focused on phytoplankton, either on specific groups, such as diatoms (Zakharova et al., 2013) or dinoflagellates (Annenkova et al., 2011), or on whole communities, from winter sub-ice waters (Bashenkhaeva et al., 2015) to spring blooms (Mikhailov et al., 2015; Mikhailov et al., 2019b; Mikhailov et al., 2019a). Remarkably few studies have aimed at charactering the diversity of all microbial eukaryotes, especially in a comparative manner. Yi et al. analyzed protist diversity by 454 sequencing of 18S rRNA gene V9-region amplicons along the Southern basin water column (52-1450 m) (Yi et al., 2017). More recently, Annenkova et al. determined the community structure of small protists (0.45-8 µm cell-size fraction) from surface waters (1-15-50 m) across the lake via 18S rRNA gene V4-region metabarcoding and suggested that some clades within known protist groups might be endemic (Annenkova et al., 2020). Nonetheless, we still lack a comprehensive view about how microbial eukaryotes distribute in the lake plankton, across basins and throughout the complete water column and, crucially, which are the most influential parameters determining community structure.

In this work, we carry out a wide-ranging comparative study of Lake Baikal planktonic protist communities in the 0.2-30 µm cell-size range using a 18S rRNA gene metabarcoding approach to study distribution patterns and to test whether depth, latitude or the coastal versus pelagic location determine community structure. With this aim, we analyze 65 samples from 17 sites across a ∼600 km latitudinal North-South transect along the three lake basins and from littoral shallow areas to deep water columns covering the epi-, meso- and bathypelagic region. Our results show complex and diverse protist communities that are mostly structured by depth and that include several typical marine lineages in low abundance. Network analyses show that epipelagic communities are much more interconnected than meso- and bathypelagic communities, suggesting potential specific biotic interactions between autotrophs, heterotrophs and parasites.

## Experimental procedures

### Sample collection

Lake Baikal water samples were collected at different depths from seventeen sites distributed along a North-South transect during a French-Russian research cruise in the summer of 2017. Sites were chosen to cover littoral (8) and open water (9) samples, including the deepest zones in the three major basins of the lake. In total, 65 water samples were collected from depths ranging from 5 to 1400 m; deep samples were collected far from the bottom to avoid sediment disturbance (Supplementary Table 1). Samples were collected with Niskin bottles (5 l for epipelagic waters, 10 l for meso- and bathypelagic waters). The physicochemical parameters of lake waters were measured with a multiparameter probe Multi 350i (WTW, Weilheim, Germany). The water was sequentially filtered onboard immediately after collection through 30-µm and 0.22-µm pore-size Nucleopore filters (Whatman, Maidstone, UK) and 0.2 µm pore-size Cell-Trap units (MEM-TEQ Ventures Ltd, Wigan, UK). Volumes of water samples filtered through Cell-Traps were smaller (samples indicated with an asterisk in Fig.1). The recovered biomass and biomass-containing filters were fixed in absolute ethanol and stored at −20°C until processed.

**Table 1.**
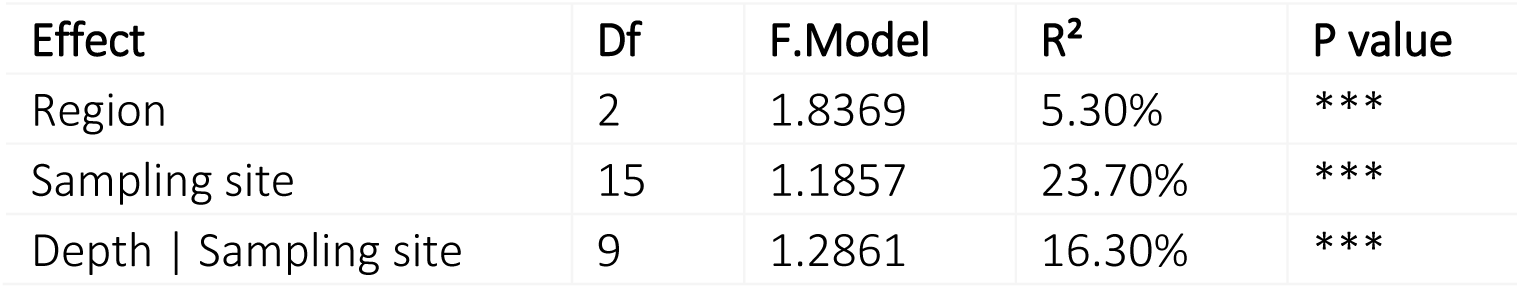
Permutational multivariate analyse of variance (PERMANOVA) of Lake Baikal planktonic protist communities across basins, sampling site and depth. PERMANOVA was calculated using Wisconsin standardization on rarefied OTUs belonging to the 65 studied plankton samples. Df, degrees of freedom.

### DNA purification, 18S rRNA gene-fragment amplification and sequencing

DNA was purified using the Power Soil™ DNA purification kit (Qiagen, Hilden, Germany). 18S rRNA gene fragments (∼530 bp) encompassing the V4 region were PCR-amplified using EK-565F-NGS (5’-GCAGTTAAAAAGCTCGTAGT-3’) and UNonMet (5’-TTTAAGTTTCAGCCTTGCG-3’), the latter biased against metazoans (Bower et al., 2004). Primers were tagged with specific 10-bp molecular identifiers (MIDs) for multiplexed sequencing. To minimize PCR-associated biases, five PCR reaction products per sample were pooled. PCR reactions were conducted in 25-µl reaction mixtures containing 0.5-3 µl of eluted DNA, 1.5 mM MgCl_2_, 0.2 mM dNTPs, 0.3 µM primers and 0.5 U Platinum Taq DNA Polymerase (Invitrogen, Carlsbad, CA) for 35 cycles (94°C for 30 s, 55-58°C for 30-45 s, 72°C for 90 s) preceded by 2 min denaturation at 94°C and followed by 5 min extension at 72°C. Pooled amplicons were purified using QIAquick PCR purification kit (Qiagen, Hilden,Germany). Amplicons were sequenced using paired-end (2×300 bp) Illumina MiSeq (Eurofins Genomics, Ebersberg, Germany). Sequences have been deposited in GenBank under the BioProject number PRJNA657482 (BioSamples SAMN15830589 to SAMN15830657).

### Sequence and phylogenetic analyses

We used an in-house bioinformatic pipeline to process raw sequences. Paired-end reads were merged with FLASH (Magoc and Salzberg, 2011) under strict criteria and assigned to specific samples based on their MIDs. MID and primer sequences were trimmed using CUTADAPT (Martin, 2011). Cleaned merged reads were next dereplicated to unique sequences using VSEARCH (Rognes et al., 2016), which was also used to detect and eliminate potential chimeras. Non-chimeric sequences from all samples were pooled together to define operational taxonomic units (OTUs) at a conservative threshold of 95% identity for 18S rRNA genes using CD-HIT-EST (Fu et al., 2012) and SWARM (Mahe et al., 2015). Singletons were excluded from subsequent analyses. OTUs were assigned to taxa based on their similarity with a local 18S rRNA database build from SILVA v128 (Quast et al., 2013) and PR2 v4.5 (Guillou et al., 2013). OTUs less than 80% identical to their best environmental hit were blasted against the GenBank *nr* database (https://ncbi.nlm.nih.gov/) and assigned manually by phylogenetic placement analyses. Briefly, the closest hits to our OTUs in SILVA and PR2 were aligned with full 18S rDNA reference sequences covering the eukaryotic diversity using MAFFT (Katoh and Standley, 2013). After removal of uninformative sites with trimAl (Capella-Gutierrez et al., 2009), we built a tree with full reference sequences with IQ-tree (Nguyen et al., 2015) under a GTR-+G+I sequence evolution model. OTU sequences were aligned to the reference alignment and then placed in the reference phylogenetic tree using EPA-ng (Barbera et al., 2019). OTUs with no reliable affiliation were maintained as ‘Unclassified’. Maximum likelihood phylogenetic trees of diplonemid and radiolarian OTUs were reconstructed from specific MAFFT alignments including their closest blast hits and reference sequences with PhyML (Guindon and Gascuel, 2003) applying a GTR+G+I (4 categories) model of sequence evolution. Bootstrap values were obtained from 100 replicates.

### Statistical analyses

We generated a table of eukaryotic OTU read abundance in the different samples of Lake Baikal for diversity and statistical analyses (Supplementary Tables 1-2). To avoid biases due to differences in absolute numbers of reads per sample, we rarefied our sequences to the second smaller number of reads (9771 in BK16.500m). BK28.100m was excluded from this process due to its lower number of reads. Statistical analyses were conducted on these data with R (R Development Core Team, 2017). Richness and diversity indices were calculated using the vegan package (Oksanen et al., 2011). Evenness was calculated according to Pielou (Pielou, 1966). To see if these indices were significantly different between sampling depths and basins, we performed Wilcoxon tests between the groups distributions using R. Likewise, to test the effect of sampling point, basin and depth class on protist community composition across samples, we conducted permutational multivariate analyses of variance (PERMANOVA) based on Wisconsin-standardized Bray-Curtis dissimilarities, using the adonis function of the vegan package. Across-sample community composition differences were visualized using non-metric multidimensional scaling (NMDS) analysis, also on Wisconsin-standardized Bray-Curtis dissimilarities. To connect communities according to specific origin we drew ellipses with the ade4 package (Dray and Dufoour, 2007). To test the significance of groups revealed by NMDS, we applied analysis of similarity (ANOSIM) tests with 999 permutations. Principal component analysis (PCA) of abiotic parameters based on centered and scaled data was performed with FactoMineR (Lê et al., 2008).

### Network analysis

We built co-occurrence networks for each depth category (epipelagic, mesopelagic, bathypelagic using a multivariation Poisson lognormal model with the PLNmodels R-package (Chiquet et al., 2018) in order to account for depth-class differences between samples and potential additional covariables (specifically the sampling basin). We retained for the analysis OTUs present in more than 20% of samples and abundances higher than 0.01%. For model selection, we used Bayesian information criteria with a 50-size grid of penalties. Networks were visualized with the ggnet R-package (Chiquet et al., 2018). To further analyze network structure, we carried out a block model analysis using a stochastic block model approach on the binary co-occurrence network using the blockmodel R-package (Leger, 2016), which synthetizes the overall network structure by gathering nodes in groups with similar modes of interactions. Network properties were calculated using the igraph R-package (Csardi and Nepusz, 2006). Properties included the number of positive and negative edges, the total number of nodes and number of connected nodes. Network mean degrees correspond to the average number of established edges. The average path length indicates the mean number of edges necessary to link a given node randomly to another. Network complexity was estimated using two indicators: connectance and clustering coefficient. The connectance was calculated as 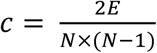, where E is the number of edges and N the number of nodes (Barrat et al., 2008). Connectance is 1 when all possible links are established. The clustering coefficient is the probability that two nodes having a similar neighbor are connected to each other (Delmas et al., 2019). It varies between 0 and 1; low values indicated poor connectivity.

## Results and discussion

### Abiotic variables across sampling sites

We collected Lake Baikal water samples along the Northern, Central and Southern basins from the same established depths in the water column (except for the deepest sample, which was collected close to the bottom but at sufficient distance –minimum 45 m– to avoid sediment influence) (Fig.1A; Supplementary Table 1). Samples from coastal areas were always collected at 5 m depth in the water column. The measured physicochemical parameters were remarkably stable across sites and depths. Temperature ranged from 3.6 to 15.3°C, but was globally low (average 5.7°C; only five surface samples exceeded 9.5°C), and significantly higher in epipelagic samples (Supplementary Fig.1). pH ranged from 7.45 to 8.47. Salinity was extremely low (always 0.0 PSU) as, accordingly, conductivity and total dissolved solids (TDS). Dissolved oxygen was high (mean 79.5%). Like temperature, pH, conductivity and dissolved oxygen in mesopelagic waters were significantly lower than in epipelagic samples.

Bathypelagic parameters were similar to those of the mesopelagic zone but more variable. In terms of basins, temperature, pH, conductivity and dissolved oxygen were higher in the Southern basin, which is also more impacted by human activities and pollution, notably aromatic hydrocarbons and mercury brought by the Selenga river (Adams et al., 2018; Roberts et al., 2020), although only oxygen and, marginally, conductivity were significantly different (Supplementary Fig.1). The two main axes of a PCA considering these abiotic parameters explained 58% of the variance (Fig.1B). Surface samples correlated with higher temperature, conductivity and, to a lower extent, pH and dissolved oxygen. These observations suggest that depth, as a proxy for light accessibility but also temperature and other abiotic parameters, might be a strong environmental driver for community structure.

### Composition of planktonic protist communities

To study the diversity and relative abundance of microbial eukaryotes in Lake Baikal plankton, we concentrated cells in the 0.2-30 µm diameter fraction by successive filtration steps. This fraction thus integrated pico-(0.2-2 µm), nano-(2-20 µm) and small microplankton (20-30 µm), covering a wider protistan spectrum than some previous comparative studies (Annenkova et al., 2020). We purified DNA and massively sequenced (MiSeq Illumina, 2×300 bp) multiplexed 18S rRNA gene V4-region amplicons. After discarding low-quality reads, we generated 6 405 343 high-quality merged paired-end sequences that we clustered in operational taxonomic units (OTUs) at different thresholds. We determined 27 504 OTUs and 9 700 OTUs at, respectively, 98% and 95% sequence identity (CD-HIT). SWARM yielded 11 590 OTUs (Supplementary Table 1), only slightly higher than the number of OTUs defined at the latter cut-off. For our subsequent comparative analyses, we deliberately retained OTUs defined at 95% sequence identity threshold. Many diversity studies focus on exact sequence variants after sequence error correction (Callahan et al., 2016) that can inform about individual strain variation. However, for the purpose of this comparative study, we chose to use conservatively defined OTUs that, on average (this varies across phylogenetic groups), correspond to the genus or species-genus level (Caron et al., 2009). This taxonomy cut-off level is relevant for broad comparative ecological studies (members of the same genus are likely to have similar general functions, despite inter-strain or species-specific niche differences), while operationally diminishing the number of handled OTUs. Based on sequence MIDs, the abundance of the different OTUs was determined for each sample (Supplementary Table 1). To avoid potential biases in diversity and relative abundance estimates linked to differences in the total number of reads, we rarefied sequences to the same number across samples, which resulted in a global number of 4 570 genus-level OTUs. Nonetheless, accumulation curves showed that the diversity of planktonic protists was far from reaching saturation, even at the conservative genus level (Supplementary Fig.2). Richness significantly decreased in deep as compared to surface waters; so did evenness (Supplementary Fig.3). A lower evenness may be partly explained by the lower cell abundance in deeper waters, as the counts for each OTU become more aleatory. We did not observe richness differences across lake basins, but evenness appeared significantly higher in the Northern basin.

From a phylogenetic perspective, our defined OTUs affiliated to at least 27 eukaryotic phyla belonging to several major eukaryotic supergroups (Fig.1C; Supplementary Table 2): the SAR clade (Stramenopiles, Alveolata, Rhizaria), Amebozoa, Archaeplastida, Excavata, Opisthokonta and Hacrobia. Although we considered Hacrobia as originally described (Okamoto et al., 2009), they should be possibly split in two or more groups as the eukaryotic phylogeny progressively resolves (Burki et al., 2020). Ciliates and dinoflagellates (Alveolata), Ochrophyta (Stramenopiles) and Holomycota (Fungi and related lineages within the Opisthokonta) dominated plankton samples representing, respectively 48.4%, 21.5%, 12.6% and 8% relative sequence abundance. Cryptophyta, Haptophyta, Kathablepharida and Tenomemida displayed moderate abundances (0.5 to 5% reads) and were followed by a long tail of lower-frequency taxa in rank:abundance curves (Supplementary Fig.4). The major dominant groups were similar in all depths, with small variations in the deepest waters. Ciliates were by far the most abundant in terms of sequence reads. However, this observation is to be pondered by the fact that, in ciliate somatic macronuclei, rRNA genes are amplified several thousand times (e.g. ∼9000 copies in *Tetrahymena thermophila* (Ward et al., 1997)), such that their relative abundance in term of cells is certainly much lower. Although diatoms (Bacillariophyta, Ochrophyta), several of them considered endemic, are well known in Lake Baikal plankton (Moore et al., 2009; Zakharova et al., 2013; Bashenkhaeva et al., 2015; Roberts et al., 2018; Mikhailov et al., 2019b), they represented only 6.1% ochrophyte reads distributed in 64 OTUs. Optical microscopy on board showed that diatoms were numerous, but their long frustules prevented most of them from being retained in the analyzed plankton fraction. Members of the Holomycota were very diverse. Classical fungi represented ca. 60% holomycotan sequences, most of them corresponding to chytrids, although the Dicarya (Ascomycota, Basidiomycota) were relatively abundant too (Supplementary Fig.5). Most Dicarya belonged to typical terrestrial fungi entering the lake waters with river in-flow or from the surrounding land. However, chytrids (flagellated fungi) are more likely to be truly planktonic organisms. Interestingly, members of Rozellida (Cryptomycota) and Aphelida, were also relatively abundant, making up to almost 40% of the holomycotan sequences. Rozellids and aphelids, together with their microsporidian relatives are parasites (Karpov et al., 2014; Bass et al., 2018). Although rozellids (cryptomycotes) are often included within fungi, they are phagotrophic organisms, unlike fungi (which are osmotrophs), and they branch more deeply than aphelids in the Holomycota tree (Torruella et al., 2018). Our data suggest that the majority of actual fungal-like planktoners in Lake Baikal are parasites.

Overall, despite methodological differences, our identified plankton protist communities were consistent with previous studies in surface waters or in a water column previously sampled in the Southern basin, with ciliates, dinoflagellates and ochrophytes being highly represented (Annenkova et al., 2020) (Yi et al., 2017).

### Marine signature taxa

Although marine-freshwater transitions are thought to be rare (Mukherjee et al., 2019) and salinity, a major driver of microbial community composition (Lozupone and Knight, 2007), high-throughput environmental studies are revealing an increasing number of typically marine eukaryotic lineages in freshwater systems. Among those are members of the parasitic perkinsids (Brate et al., 2010), haptophytes (Simon et al., 2013), Bolidophyceae (Richards and Bass, 2005; Annenkova et al., 2020) and several Marine Stramenopiles (MAST) clades (Massana et al., 2004; Massana et al., 2006), such as MAST-2, MAST-12, MAST-3 and possibly MAST-6 (Simon et al., 2015a). Recently, diplonemids, a cosmopolitan group of oceanic excavates particularly abundant and diverse in the deep ocean (Lara et al., 2009; de Vargas et al., 2015) were identified in deep freshwater lakes (Yi et al., 2017; Mukherjee et al., 2019). Likewise, Syndiniales, a clade of parasitic alveolates (often parasitizing their dinoflagellate relatives) widely distributed in oceans (López-García et al., 2001; Guillou et al., 2008), were recently identified in Baikal surface plankton (Annenkova et al., 2020). We identified members of all these lineages in our large Lake Baikal plankton dataset, albeit mostly in low proportions (Fig.2A; Supplementary Table 3). Bolidophytes and, collectively, MAST clades were nonetheless relatively abundant in the lake. However, MAST clades are not monophyletic and they exhibited different abundance patterns. Clades previously detected in freshwater systems, MAST-2, MAST-6, MAST-12 and to a lesser extent MAST-3, were relatively abundant. But MAST clades not previously observed in other freshwater systems, including MAST-1, MAST-4, MAST-8 and MAST-20 occurred in very low proportions in a few samples. In addition to the rare diplonemids, which were widely but sporadically present across Lake Baikal samples (Fig. 2A-B), we identified OTUs belonging to the emblematic Radiolaria, to our knowledge never before identified in freshwater plankton. These OTUs were members of the Polycystinea (Fig.2C) and exhibited extremely low frequencies.

**Fig. 1.**
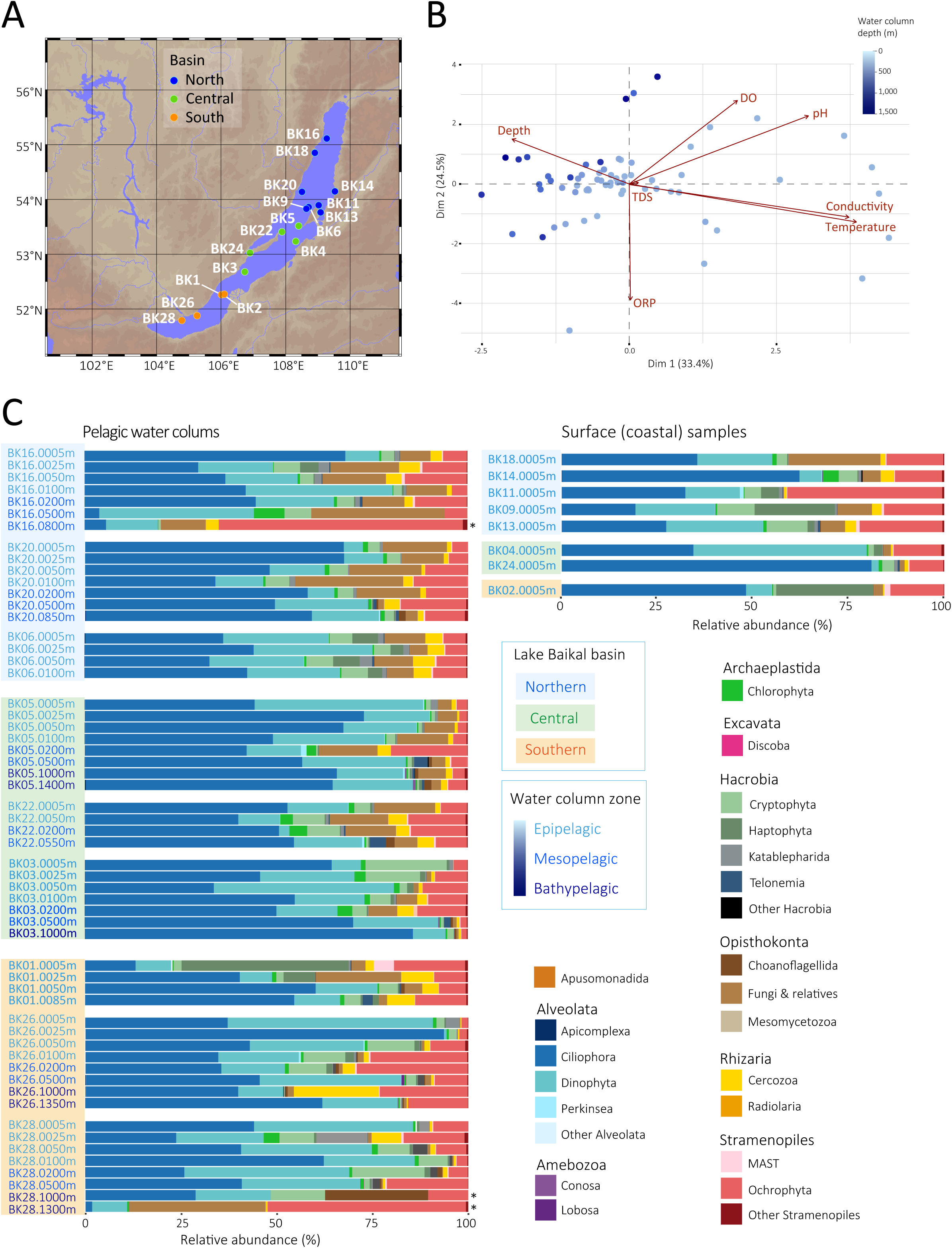
Sampling sites and overall planktonic protist community composition in Lake Baikal. **A**, map of Lake Baikal showing the different sampling sites across the three major lake basins (indicated by colors). **B**, Principal component analysis (PCA) of samples according to their associated physicochemical parameters. The number near the points correspond to the sampling site, and the color of the points indicates their sampling depth. TDS, total dissolved solids; DO, dissolved oxygen; ORP, oxidation-reduction potential. Blue tones indicate the sampling depth in the water column, as indicated. **C**, Relative abundance of different high-rank eukaryotic taxa in Baikal plankton based on read counts for the defined OTUs. The asterisk indicates samples retrieved from Cell-Traps (Methods). Color codes for sample basin and depth origin as well as for the different taxa are indicated.

**Fig. 2.**
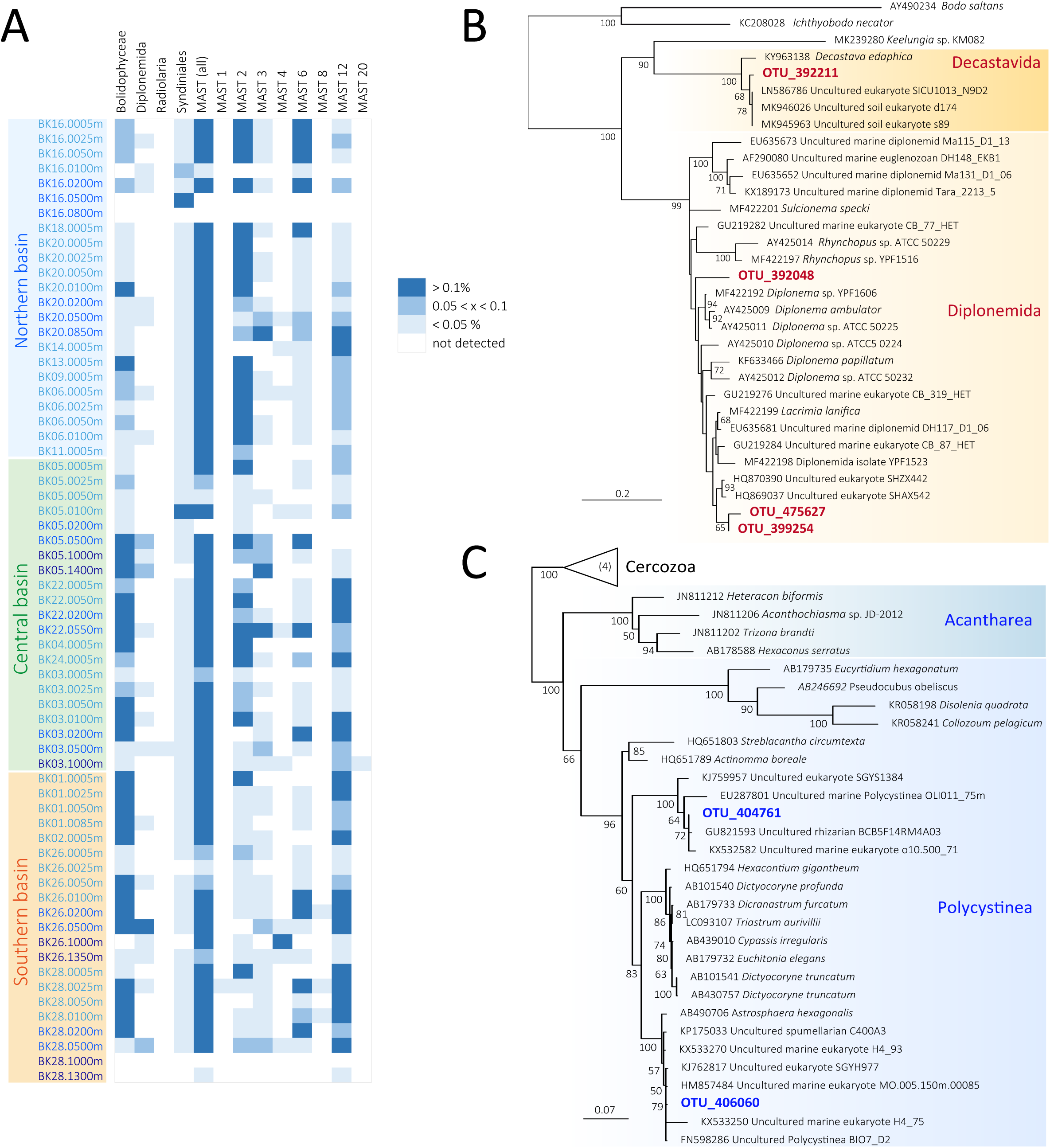
Marine signature taxa detected in Lake Baikal plankton. **A**, heat map showing the relative abundance of different typically marine taxa across Baikal plankton samples. The frequency of the different phylogenetic groups is indicated by different shades of blue. **B**, Maximum Likelihood (ML) phylogenetic tree of OTUs belonging to diplonemids and a related group of euglenozoan excavates (594 unambiguously aligned positions). **C**, ML tree of radiolarian OTUs (534 unambiguously aligned positions).

The low abundance of some of these typically marine lineages partly explains the fact that they failed to be detected in previous studies of freshwater systems, suggesting that these ecological transitions have been so far underestimated (Paver et al., 2018). However, an additional explanation might be found in the particular features of the Lake Baikal, including its considerable depth, marked oligotrophy and even the presence of deep-venting (Müller et al., 2001; Sherstyankin et al., 2006; UNDP-GEF, 2015), which make it qualify in all points but salinity as a freshwater sea.

### Environmental drivers of protist community structure

To test whether planktonic protist communities were influenced by abiotic factors (clearly correlated to sample spatial origin; Fig.1), we carried out permutational multivariate analysis of variance (PERMANOVA) of Wisconsin-standardized Bray-Curtis distances between communities as a function of sample spatial origin. PERMANOVA tests revealed significant differences in microbial eukaryotic communities as a function of basin (latitudinal region), sampling site (coordinates) and depth within sampling sites (Table 1). However, the most influential effects were those of the water column location, 23.7%, which combine latitudinal and vertical determinants, and depth within each single water column, i.e. vertical variation alone (16.3%). The effect of the sampling basin was significant but small (5.3%). To better visualize differences between communities, we carried out an NMDS analysis on the global Bray-Curtis distance matrix. Points from most water columns did not show a marked differentiation, as most water columns overlapped to some extent (Supplementary Fig.6). Likewise, samples from different basins did not show a clear differentiation, although samples from the Southern basin tended to segregate from the two other basins (Fig.3A). Samples from coastal versus open waters did not segregate at all (Fig.3B). However, planktonic communities clearly segregated as a function of the water column zonation, with epipelagic, mesopelagic and bathypelagic communities well separated in the NMDS plot (Fig.3C). NMDS analyses based on SWARM-defined OTUs yielded very similar results (Supplementary Fig.7). These observations were statistically supported by ANOSIM tests, which showed significant and marked differences among communities according to depth, significant but weak differences according to basin origin, and no correlation at all between coastal and pelagic samples (Supplementary Table 4).

These results suggest that depth is the major environmental factor structuring Lake Baikal protist communities. Depth is in turn a proxy for a variety of abiotic parameters, notably light, but also, despite their limited variation, temperature, dissolved oxygen, conductivity and pH (Fig.1). These environmental variables and others, such as the nature of dissolved organic matter (TDS amount does not vary significantly; Fig.1B), are likely to influence prokaryotic communities as well (Kurilkina et al., 2016). Consequently, the nature of prey available for bacterivorous protists is possibly different. This may, in turn, select for protists with particular preying affinities, such that biotic interactions with other planktonic members may be also important determinant factors of community structure and function.

### Functional groups and biotic interactions

To look for potential ecological interactions between members of protist communities, we first explored the distribution of major functional classes with depth. We attributed protists to three major categories based on knowledge about the lifestyle and ecological function of the corresponding phylogenetic lineages: autotrophs, free-living heterotrophs and parasites (Supplementary Table 5). We acknowledge that these are very broad categories and that many photosynthetic organisms can be mixotrophs (Massana, 2011; Mitra et al., 2016). However, information about mixotrophy is still scarce and it is difficult to predict this ability from sequence data only. Therefore, our category ‘autotrophs’ included also photosynthetic organisms that can additionally use heterotrophic feeding modes. Free-living heterotrophs include predatory protists but also osmotrophic organisms feeding on organic matter, such as fungi or some Stramenopiles. The relative abundance of the three functional categories in Lake Baikal significantly followed the same trend in the three water column zones, with autotrophs being less abundant than heterotrophs and parasites being in much lower proportion (Fig.4). Low proportions of parasitic protists are consistent with affordable parasite loads for an ecosystem, as was previously observed (Simon et al., 2015b). Nonetheless, the relative amount of parasitic lineages diminished with depth, potentially suggesting that a relatively important proportion of protists identified in deep waters might be inactive. This is indeed likely the case for most photosynthetic organisms that were identified below the epipelagic region. Although the proportion of autotrophs diminished with depth, they still made up to 30% of the total in bathypelagic waters. As mentioned, some of these protists may be mixotrophic and prey on bacteria or other protists in the dark water column. However, the majority of photosynthetic lineages may simply be inactive, dormant or on their way to decay, serving as food for the heterotrophic component of microbial communities. The presence of relatively abundant photosynthetic protists in the Baikal dark water column and sediments is well documented (Zakharova et al., 2013; Yi et al., 2017), low temperatures possibly helping their preservation during sedimentation. Finally, free-living heterotrophs were the most abundant functional category throughout the water column. This might seem at odds with a pyramidal food-web structure whereby primary producers should be more abundant than consumers. However, several factors might explain this. First, the presence of ciliates likely introduces a positive bias in this functional category. Second, many autotrophs might be, on average, larger than heterotrophic protists and their biomass exceed that of consumers. Finally, many heterotrophic protists might depend on bacteria or on larger organisms (e.g. fungi degrading decaying plant material).

**Fig. 3.**
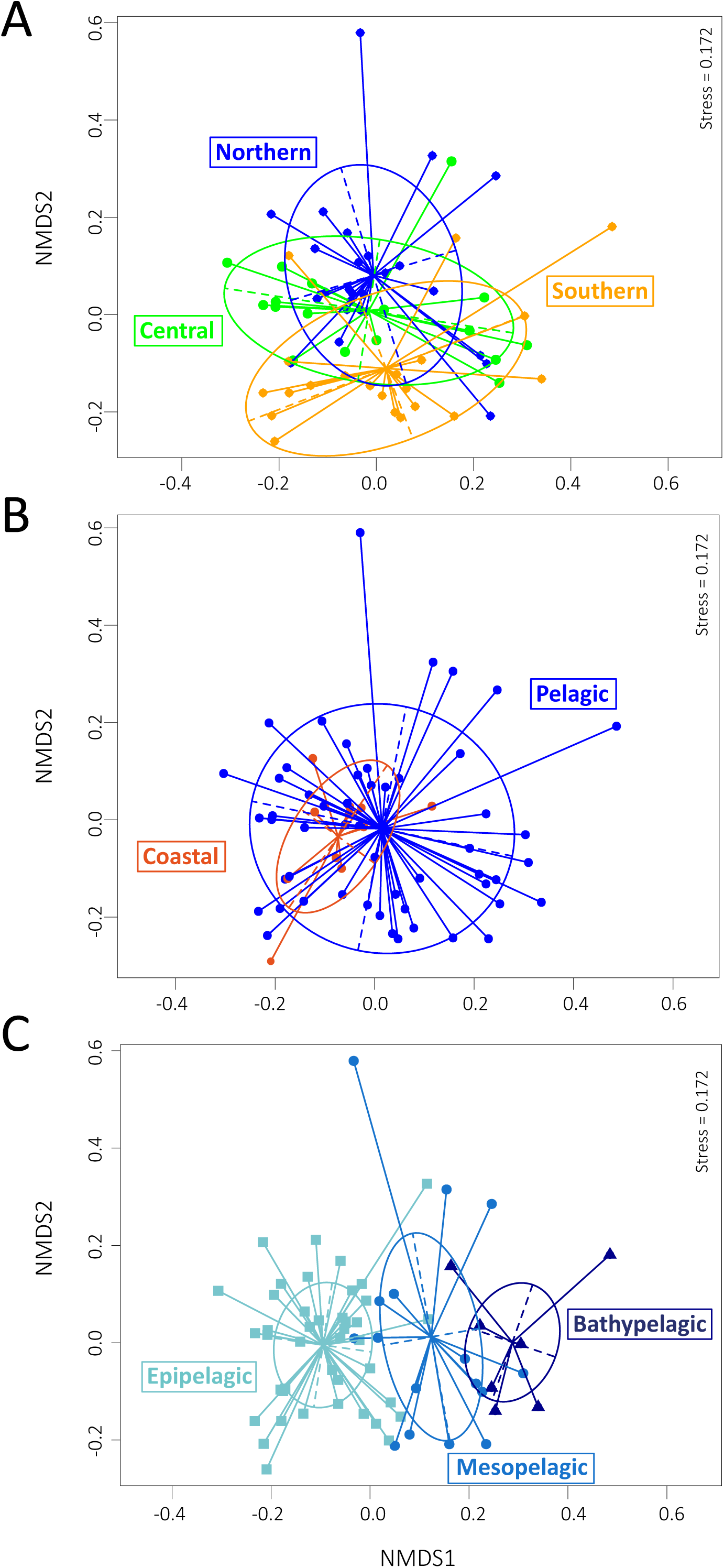
Non-metric multidimensional scaling (NMDS) analysis of Lake Baikal plankton samples as a function of protist community similarities. The NMDS plot was constructed with Wisconsin-standardized Bray-Curtis dissimilarities between all samples. A, plankton samples highlighted by basin origin. B, plankton samples from coastal, shallow sites versus open water sites. C, samples grouped according to their depth origin in the water column; epipelagic (<200 m), mesopelagic (200-500 m), bathypelagic (>500 m).

**Fig. 4.**
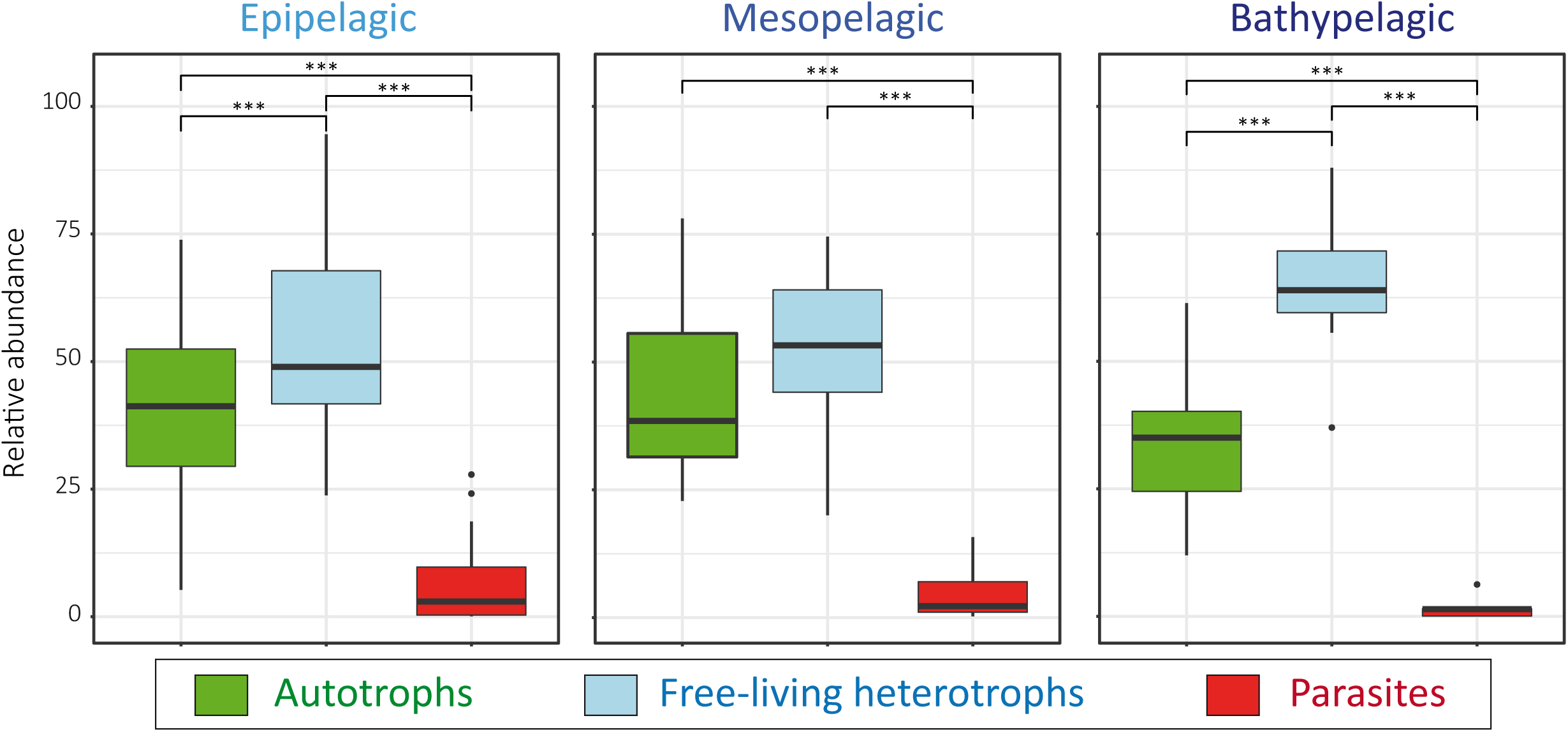
Box plots showing the distribution of relative abundances of major functional categories of protists in Lake Baikal plankton. The three plots show the relative abundance of sequences affiliated to autotrophic, heterotrophic and parasitic protists for each sampling depth class. The thickest line inside each box represents the median on the distribution, bottom and top borders of boxes correspond to the first and the third quartiles and whiskers extend to minimal and maximal distances. Significant differences between distributions are indicated with stars (p-values <0.05, <0.005 and <0.0005 are respectively indicated by one, two and three stars). For the assignation of taxa to functional categories, see Supplementary Table 5.

To further explore biotic interactions, and given that protist community differences were essentially seen throughout the water column, we reconstructed co-occurrence networks of OTUs found in epipelagic, mesopelagic and bathypelagic zones. To build the networks, we retained OTUs present in more than 20% of samples at relative abundances higher than 0.01% (Supplementary Table 6). The structure of the three networks was markedly different (Fig.5). The epipelagic network was denser, having more interconnected OTUs, more positive interactions and several hub-type OTUs that interact with many OTUs. Mean node degrees were also higher in the epipelagic network (Supplementary Table 7). Meso- and bathypelagic networks had less connected nodes and most correlations were negative. Only one positive interaction was observed in mesopelagic waters (ciliate-fungus) and in bathypelagic waters (rozellid-ochrophyte). The latter might suggest a specific parasitic interaction. Although bathypelagic waters exhibited the least connected nodes, both the connectance and the clustering coefficient of the network were the highest. A block-model representation of the three networks indicated the occurrence of pairs of OTU sets sharing similar properties that were highly interconnected with each other and only loosely to other sets (Supplementary Fig.8).

**Fig. 5.**
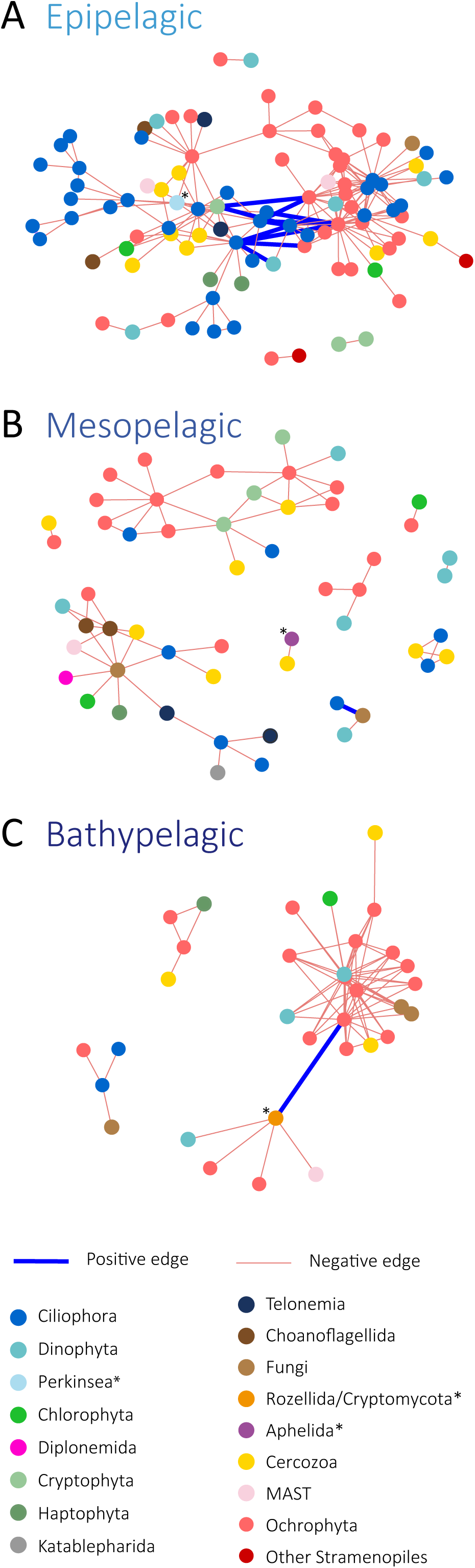
Co-occurrence networks of planktonic protists in the Lake Baikal water column. A, network obtained from epipelagic (<200 m) samples across the lake. B, network obtained from mesopelagic (200-500 m) samples. C, network obtained from bathypelagic (>500 m samples). Networks were built on OTUs present in more than 20% samples and having a relative abundance higher than 0.01%. OTUs are represented by nodes and direct covariations between them, by edges. Nodes and taxa labeled with an asterisk correspond to putative parasites.

Collectively, our network data suggest that epipelagic communities are more active and have more positive and negative interactions, whereas in very deep waters, communities are more stable with a restricted but strongly connected core.

### Concluding remarks

Lake Baikal in Southern Siberia is a unique freshwater system by its volume, maximum depth (1 642 m) and topographical features that include rifting associated with hydrothermalism. With its highly oligotrophic waters, it amounts to an inner freshwater sea in all points but an extremely low salinity. Freshwater ecosystems are particularly threatened by climate change and, being located in Southern Siberia, one of the most rapidly changing zones, Lake Baikal is being severely impacted (Mackay et al., 2017). Yet, despite the importance of the lake and its uniqueness, its microbial planktonic communities have been only partially studied and we lack reference comprehensive comparative community data to assess ongoing and future change and infer how it may affect microbial functions and the ecology of the lake. In this study, we have analyzed the composition of microbial eukaryotic communities in plankton collected from different water columns along a transect of ∼600 km across the three lake basins, from surface (5 m) to high depth (1 400 m) and from littoral to open waters. Our study shows widely diverse communities covering all eukaryotic supergroups, with ciliates, dinoflagellates, chrysophytes and flagellated fungi plus related lineages (rozellids, aphelids) being the most relatively abundant, together with cryptophytes, haptophytes, katablepharids, telonemids and several MAST lineages. Interestingly, confirming previous observations in Lake Baikal, we observed members of typically marine lineages, including bolidophytes, syndineans, diplonemids and, for the first time, radiolarians. These observations suggest that the salinity barrier is relatively easy to cross and that the ‘marine’ determinants might be more related with the oligotrophic nature of the system and the occurrence of a deep water column than with salinity itself. Despite the relatively homogeneous values of several physicochemical parameters, planktonic protist communities were highly and significantly stratified in Lake Baikal, suggesting that depth, as a proxy for light but also temperature, pH, oxygen and conductivity, is a major determinant of community structure. By contrast, the effect of latitude (basins) was minor, if not negligible. Consistent with vertical stratification, the relative proportion of autotrophs, free-living heterotrophs and parasites is altered with depth, where photosynthetic lineages are still present but, like parasites, in lower proportions. Biotic factors are also important in structuring Lake Baikal communities. Co-occurrence network analyses showed highly interconnected communities in surface waters, with positive and negative interactions. By contrast, deep, bathypelagic communities exhibit much less connected OTUs, although they are strongly, and mostly negatively, correlated. This might be suggestive of much more diluted and potentially inactive populations, but with a conserved core of highly interconnected OTUs. Our results pave the way for future comparative analyses of protist communities through time, notably in the context of rapid climate change that is affecting Siberia and Lake Baikal.

## Acknowledgments

We thank the crew of the R/V G. Titov for their professionalism and efficiency onboard, the director of the Limnological Institute at Irkusk for logistical assistance and Philippe Deschamps for technical bioinformatic support. This research was funded by the European Research Council Grants ProtistWorld (322669, PL-G) and PlastEvol (787904, DM) as well as the Russian State grant 0345-2016-0009 (NVA).

## Author contributions

PLG, DM and NVA designed the work and organized the limnological cruise. PLG, PB, AILA, LG, GR and NVA collected and processed water samples during the cruise. PB purified DNA and carried out PCR reactions for metabarcoding analysis. GR carried out the initial bioinformatic analysis of amplicon sequences. GD carried out metabarcoding, statistical and network analyses, with help from LJ. PLG wrote the manuscript with input from co-authors. All authors read, critically commented and approved the final manuscript.

