## Supplementary material for "Environmental drivers of plankton protist communities along latitudinal and vertical gradients in the oldest and deepest freshwater lake"

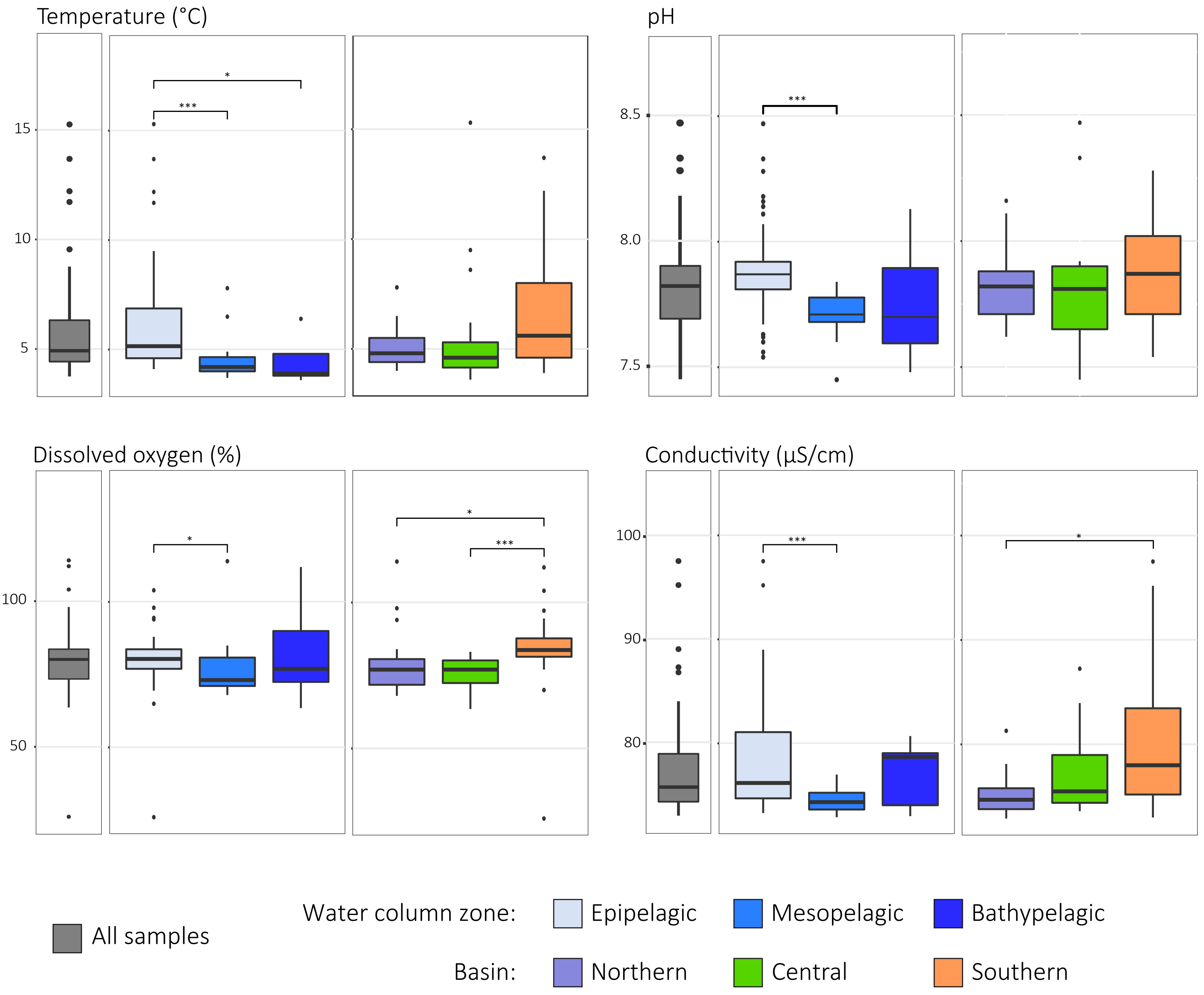

**Supplementary Fig.1.** Box plots of physico-chemical parameters measured in Lake Baikal water samples. Average values and variation are shown for all the Baikal samples and as a function of basin and depth. Significant differences between pairs of samples are indicated (p-value ≤0.05*; p-value <0.001***).

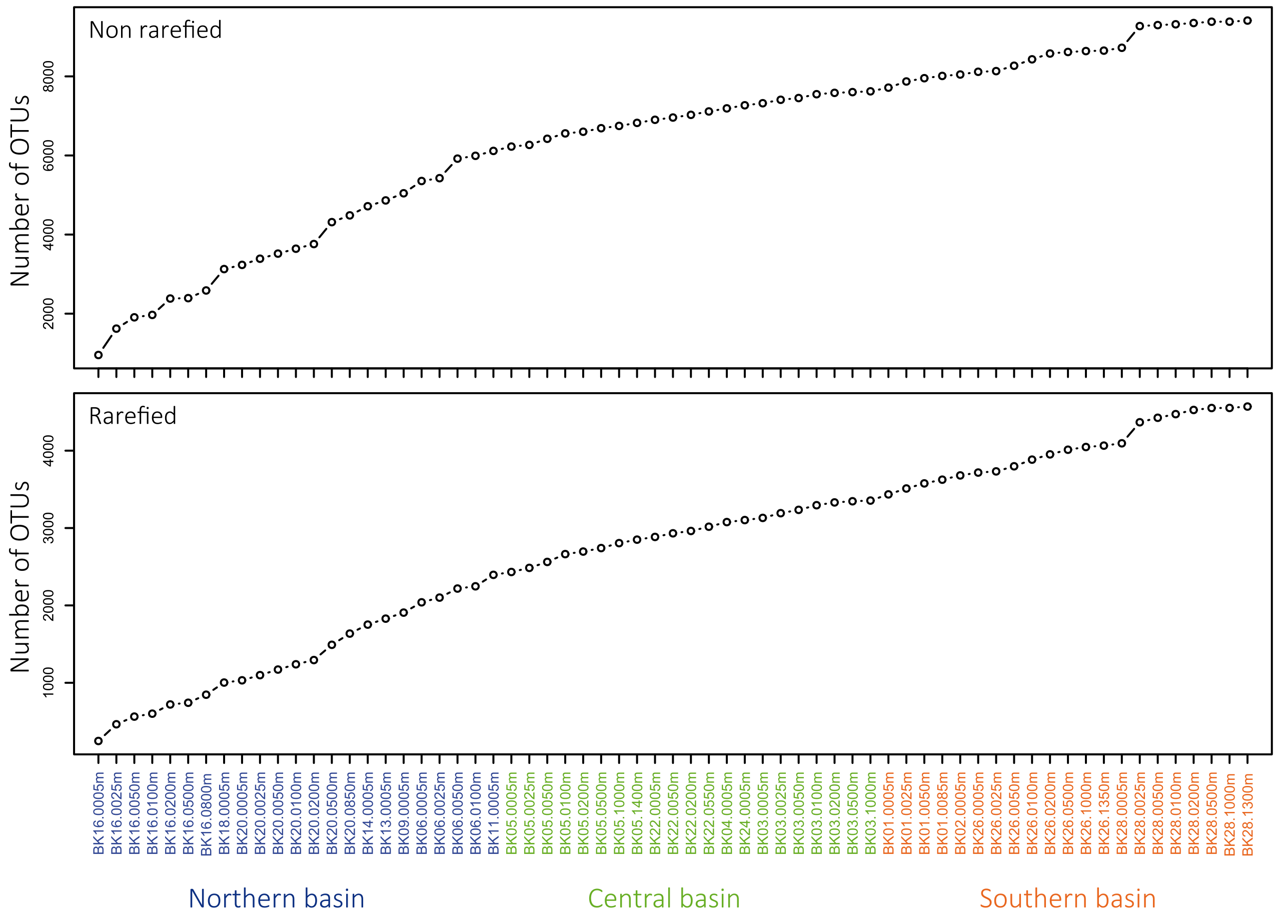

**Supplementary Fig. 2.** Accumulation curves for OTUs identified in Lake Baikal plankton samples before and after rarefaction.

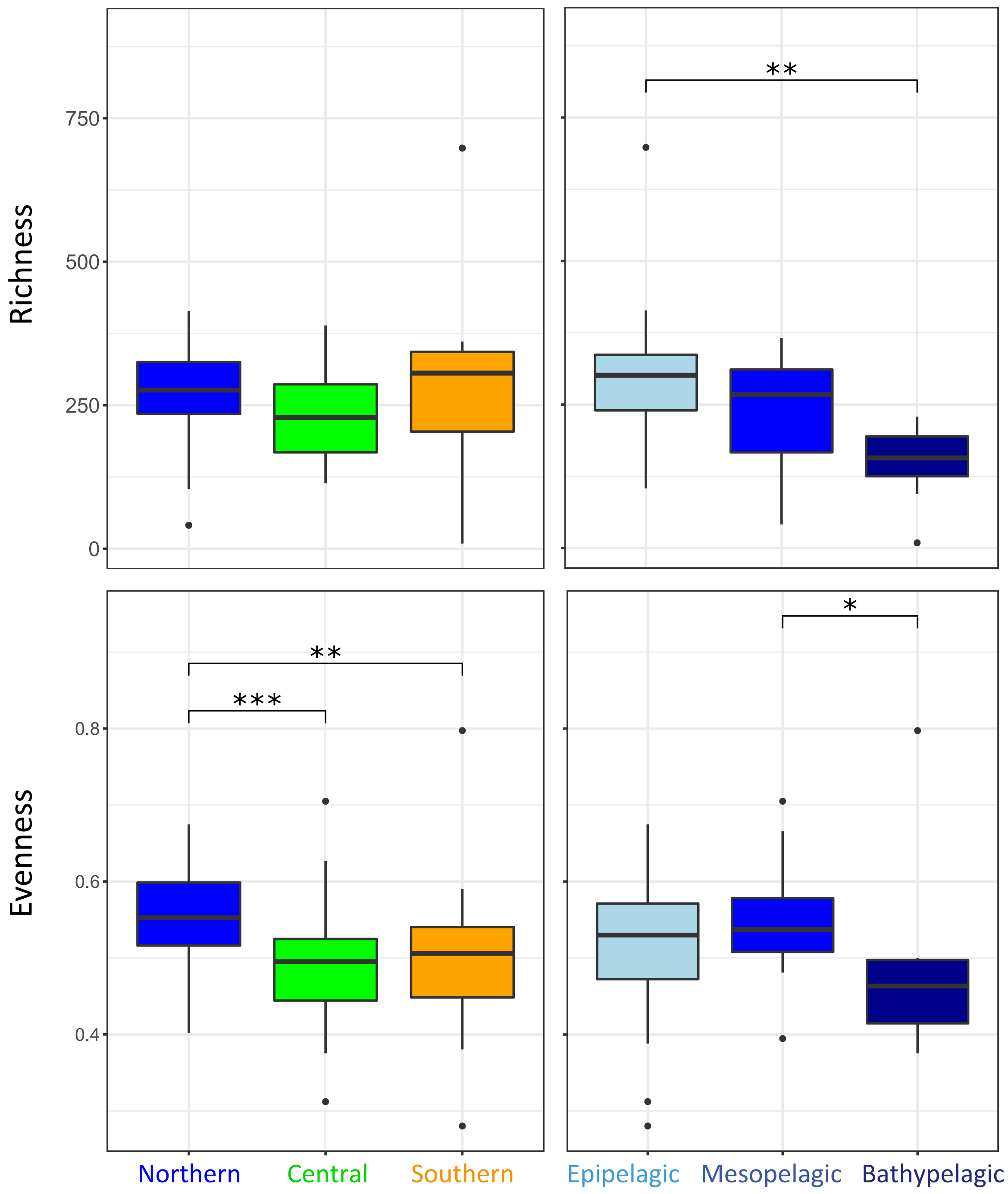

**Supplementary Fig.3.** Box plots showing diversity (richness) and evenness values in Lake Baikal water samples as a function of basin and depth. Richness and evenness were calculated on OTUs defined at 95% 18S rRNA gene sequence identity (~genus level). Significant differences between pairs of samples are indicated (p-value ≤0.05*; p-value <0.001***).

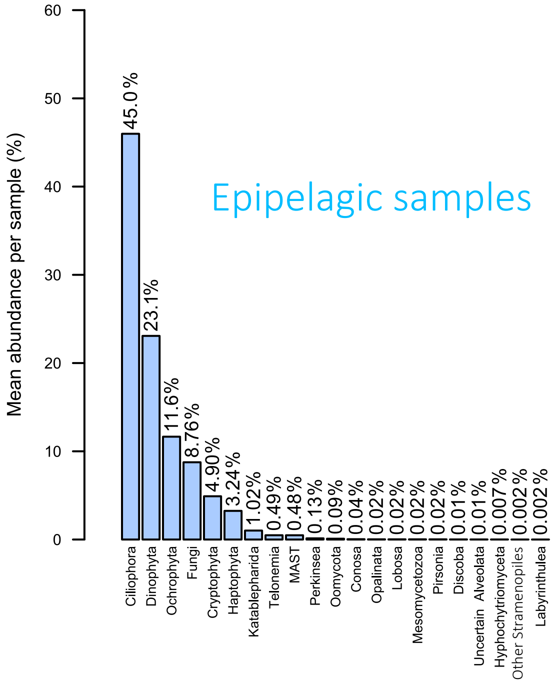

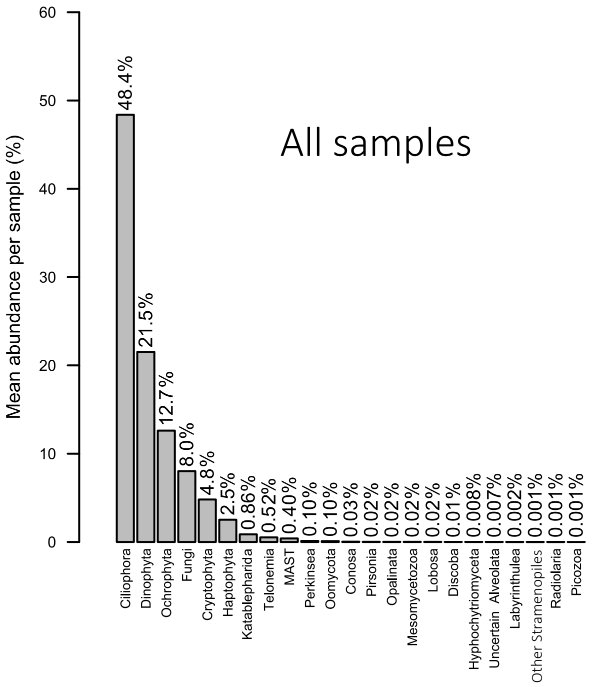

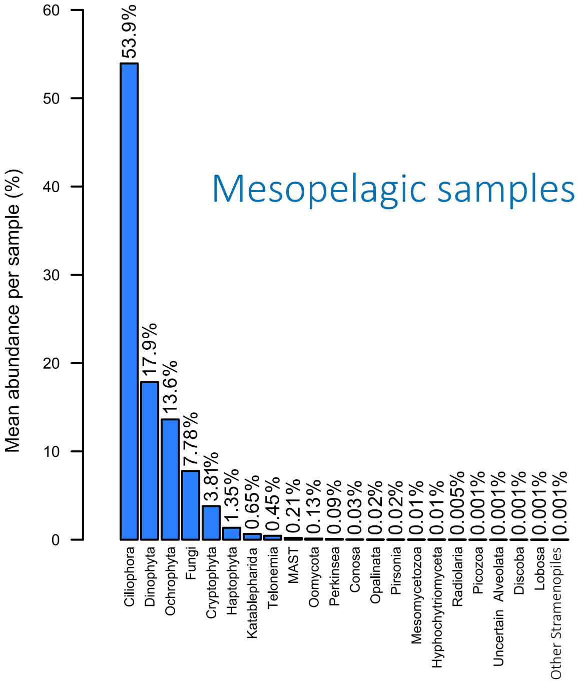

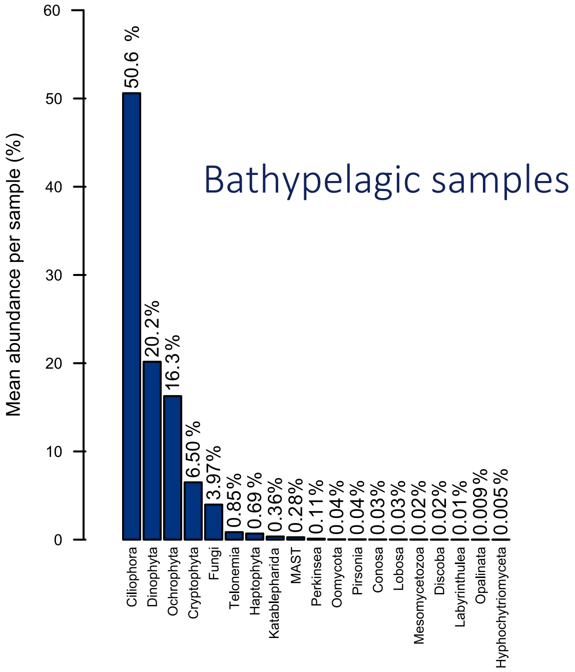

**Supplementary Figure 4.** Rank:abundance curves of protist OTUs in Lake Baikal plankton samples. Rank:abundance curves are presented globally for the lake and by lake depth category.

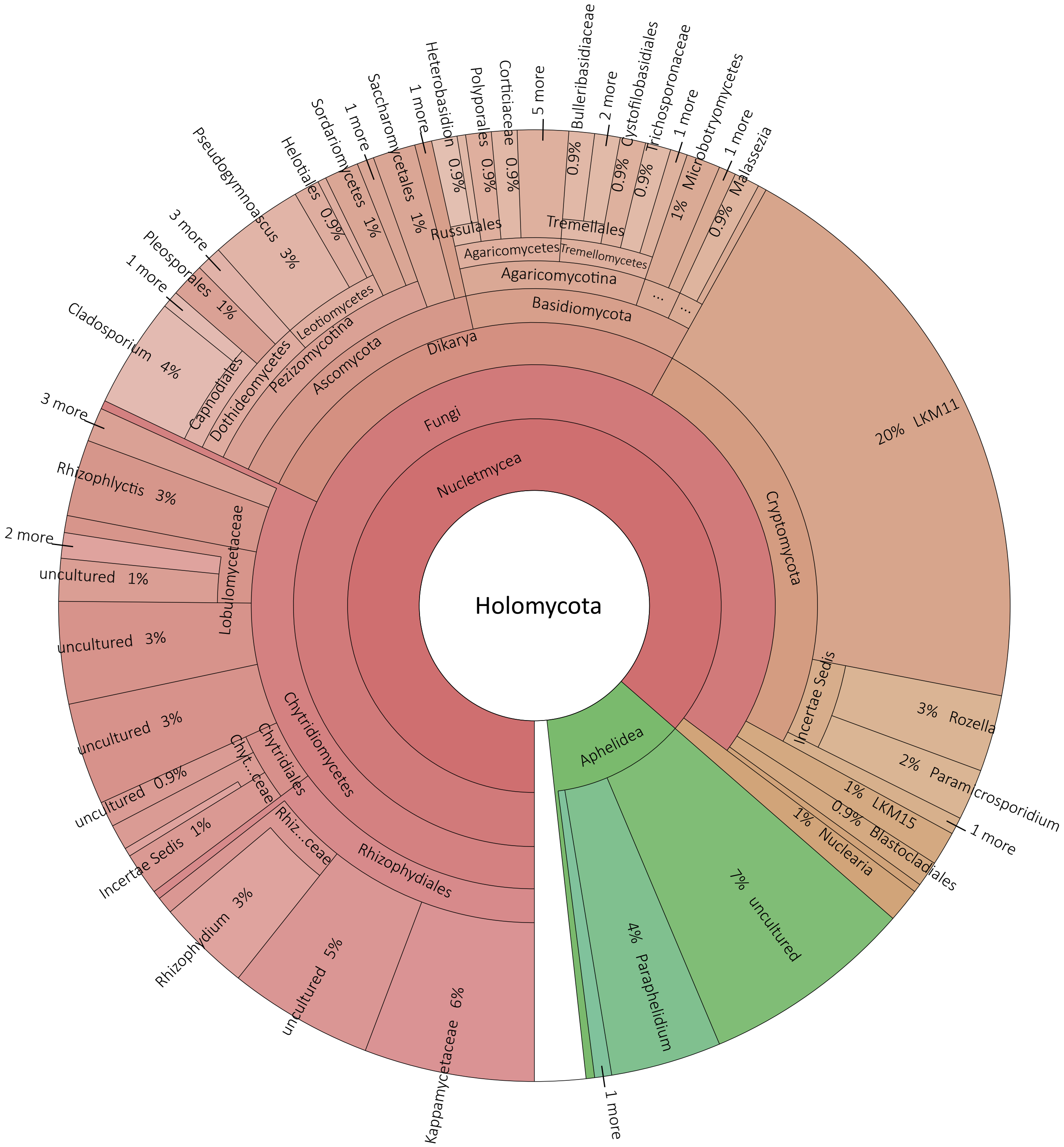

**Supplementary Figure 5.** KRONA representation of the global diversity of Holomycota in Lake Baikal plankton. Holomycota is one of the two branches of Opisthokonta, including Fungi and related lineages (Cryptomycota, Aphelida, Nuclearida). The graph was produced using the SILVA classifier (https://www.arb-silva.de).

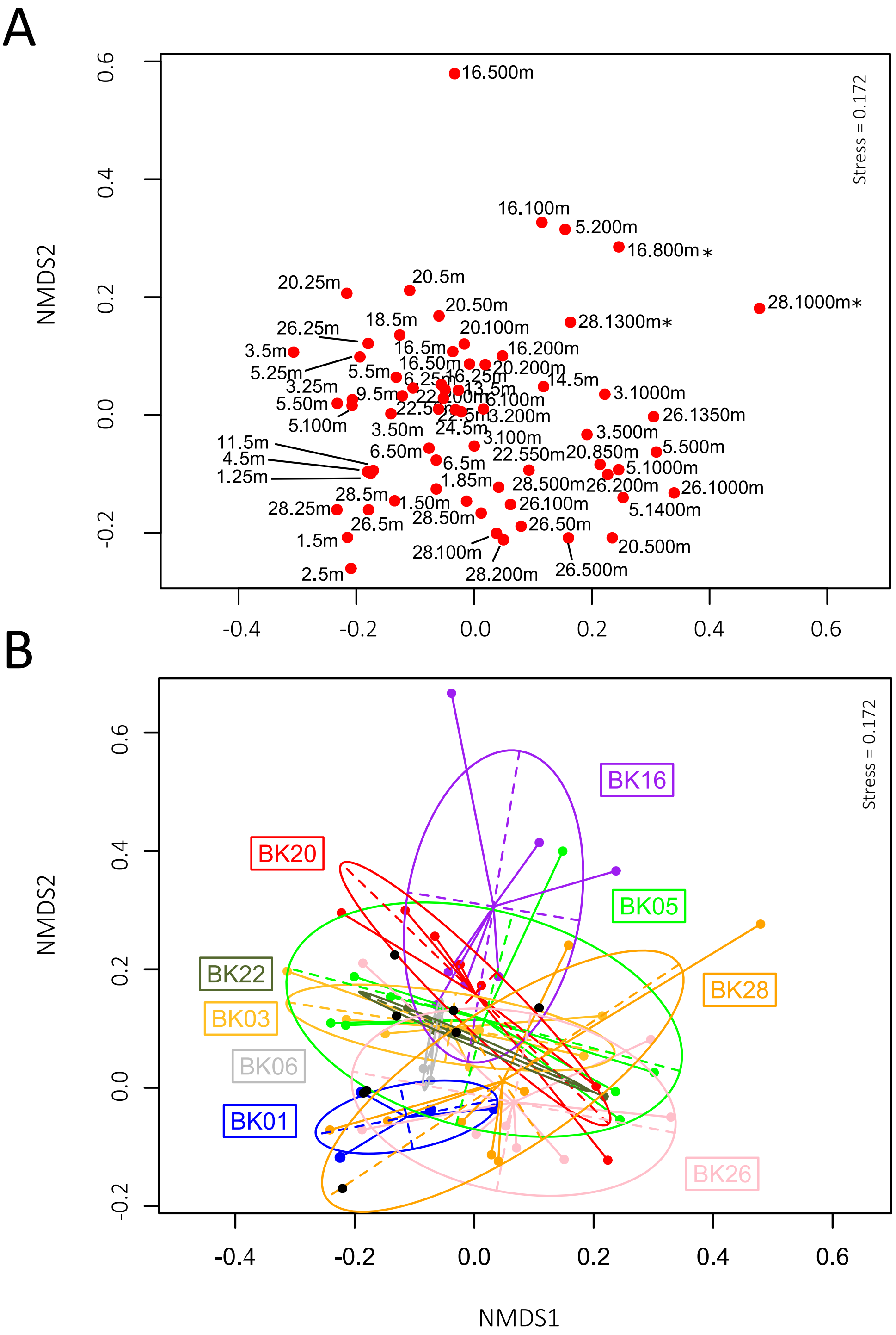

**Supplementary Fig. 6.** Non-metric multidimensional scaling (NMDS) analysis of Lake Baikal plankton samples based on differences in protist community composition. Community structure differences are based on Bray-Curtis distances established upon the distribution and relative frequency of OTUs. A, NMDS showing all sampling points. B, NMDS on the same points taking into account different water column samples. Black dots correspond to surface (5 m deep), coastal samples.

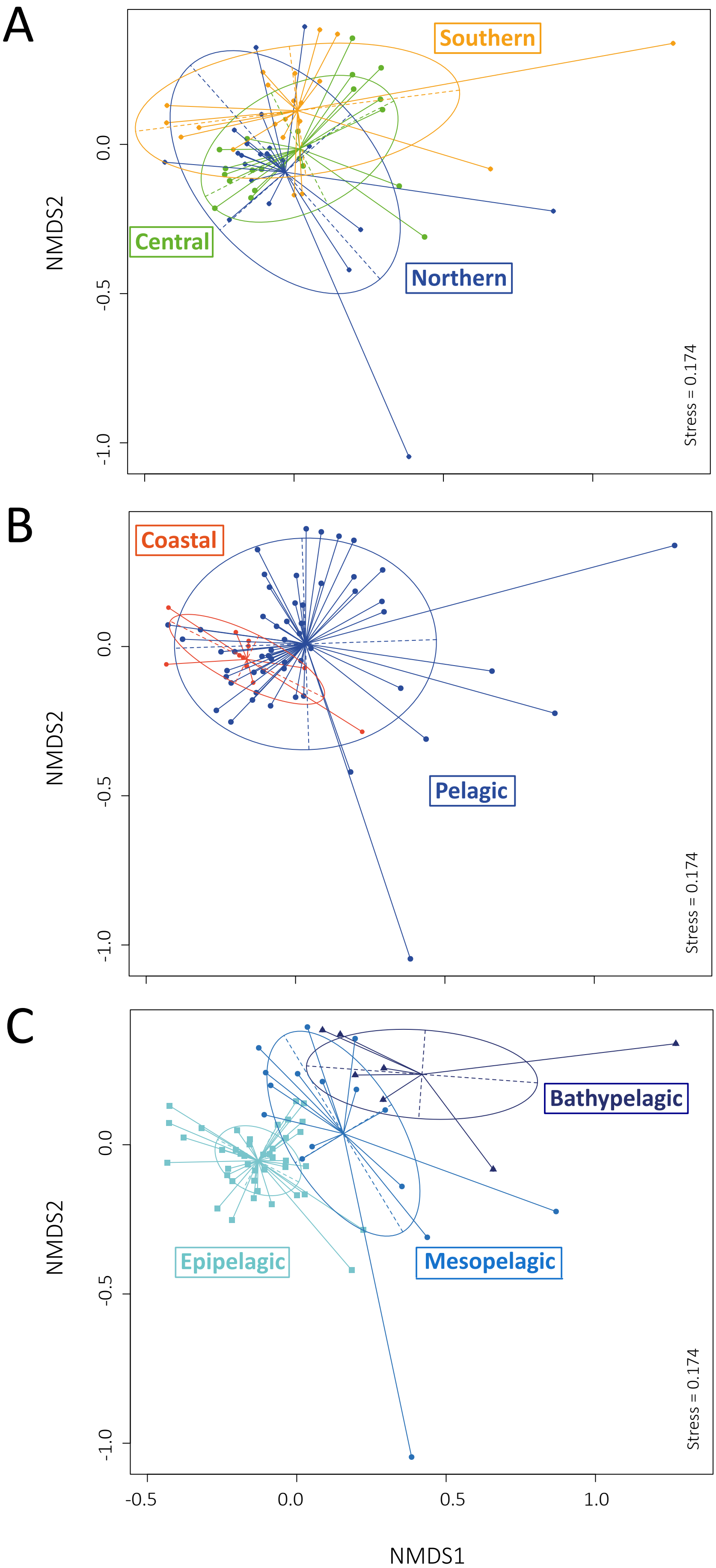

**Supplementary Fig. 7.** Non-metric multidimensional scaling (NMDS) analysis of Lake Baikal plankton samples as a function of protist community similarities based on SWARM-determined OTUs. The NMDS plot was constructed with Wisconsin-standardized Bray-Curtis dissimilarities between all samples. A, plankton samples highlighted by basin origin. B, plankton samples from coastal, shallow sites versus open water sites. C, samples grouped according to their depth origin in the water column; epipelagic (<200 m), mesopelagic (200-500 m), bathypelagic (>500 m).

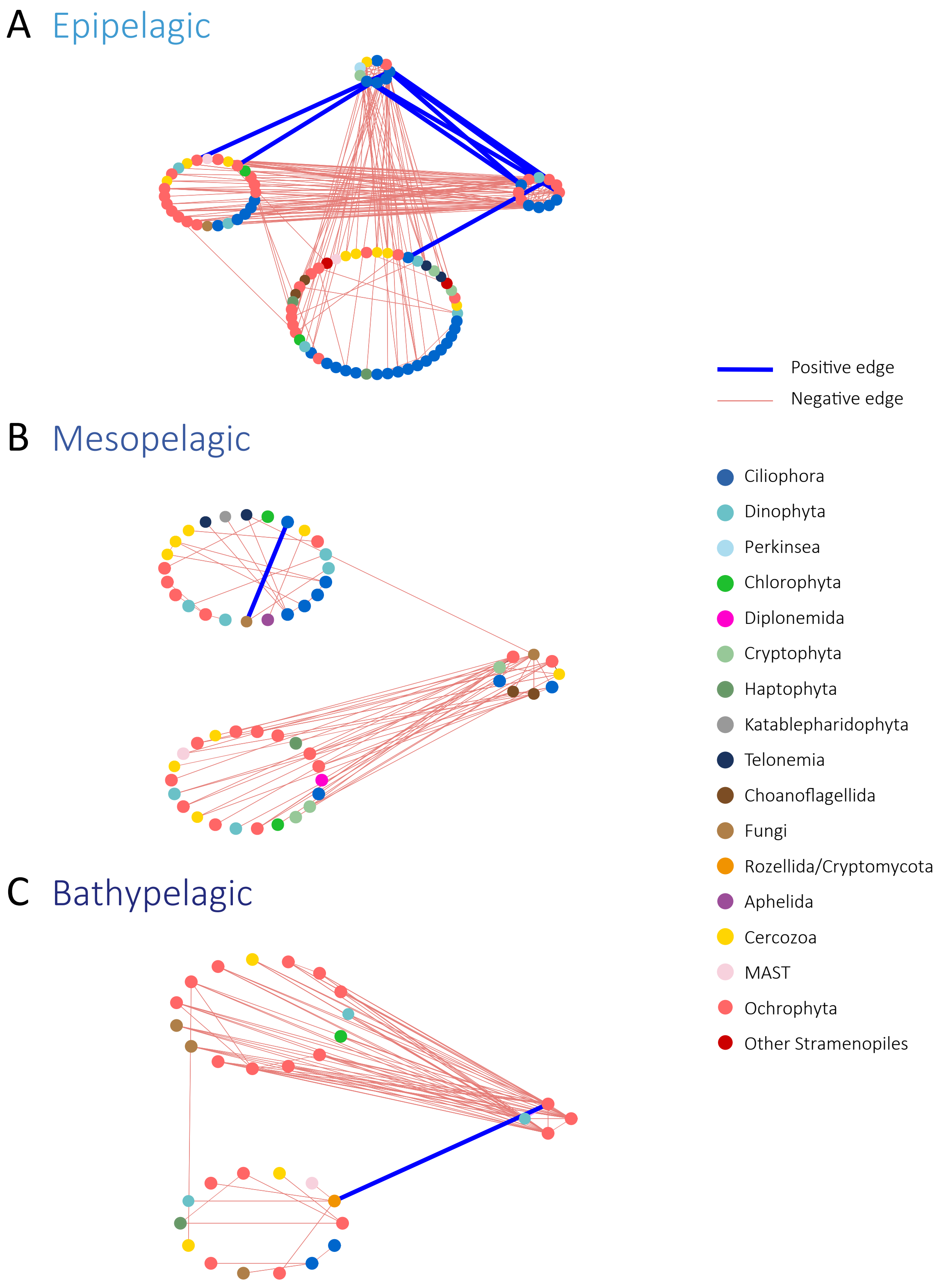

**Supplementary Figure 8.** Block-model representation of the networks of the planktonic protists in Lake Baikal for the three major depth categories. Networks were built on OTUs present in more than 20% samples and having a relative abundance higher than 0.01%. OTUs are represented by nodes and direct covariations between them, by edges.

**Supplementary Table 1.** Sampling sites in Lake Baikal water column and associated physico-chemical, sequence and diversity data. Operational taxonomic units (OTUs) were defined at 95% sequence identity, except when indicated (SWARM). Singletons are excluded from the OTU counts.

| Basin | Sampling site | Sample name | Sampling depth (m) | Sampling Date | Latitude | Longitude | Bottom depth (m) | Temperature (°C) | pH | Conductivity (µS/cm) | TDS (mg/l) | ORP (mV) | DO (%) | Raw reads | Clean merged reads | Number of OTUs | Number of SWARM OTUs | No. OTUs after rarefaction | Simpson | Eveness |
|---|---|---|---|---|---|---|---|---|---|---|---|---|---|---|---|---|---|---|---|---|
| North | BK16 | BK16.5m | 5 | 02.07.2017 | 55°06'259" N | 109°16'104" E | 846 | 4.7 | 7.88 | 75.7 | 73.5 | 106.7 | 76 | 176014 | 132046 | 954 | 1216 | 248 | 0.87 | 0.51 |
| | | BK16.25m | 25 | | | | | 4.6 | 7.67 | 75.7 | 69.6 | 79.5 | 69.5 | 129314 | 103211 | 1156 | 1128 | 357 | 0.94 | 0.6 |
| | | BK16.50m | 50 | | | | | 4.4 | 7.82 | 74.3 | 73 | 106.1 | 80.5 | 109882 | 91554 | 854 | 1052 | 276 | 0.94 | 0.6 |
| | | BK16.100m | 100 | | | | | 4.3 | 8.16 | 74.7 | 70.9 | 106.1 | 71.8 | 70538 | 68416 | 197 | 167 | 104 | 0.85 | 0.55 |
| | | BK16.200m | 200 | | | | | 4.2 | 7.81 | 74.4 | 74.1 | 105.5 | 73 | 150492 | 137629 | 980 | 1309 | 286 | 0.91 | 0.56 |
| | | BK16.500m | 500 | | | | | 4.2 | 7.68 | 74.2 | 77.4 | 101.8 | 68 | 10085 | 9771 | 41 | 26 | 41 | 0.89 | 0.67 |
| | | BK16.800m.CT | 800 | | | | | 4.2 | 7.68 | 73.8 | 76.1 | 67.6 | 71.8 | 56767 | 51846 | 261 | 54 | 135 | 0.82 | 0.52 |
| | BK18 | BK18.5m | 5 | 03.07.2017 | 54°51'372" N | 108°54'14" E | 36 | 5.6 | 8.07 | 77.5 | 76.1 | 100.3 | 94 | 136920 | 95373 | 1178 | 1039 | 322 | 0.91 | 0.54 |
| | BK20 | BK20.5m | 5 | 03.07.2017 | 54°08'361" N | 108°30'613" E | 896 | 5.6 | 7.92 | 73.9 | 74.8 | 136.1 | 75.8 | 130943 | 102135 | 518 | 712 | 147 | 0.7 | 0.4 |
| | | BK20.25m | 25 | | | | | 4.9 | 7.9 | 73.6 | 76.1 | 134.8 | 78 | 149603 | 102765 | 698 | 927 | 203 | 0.73 | 0.42 |
| | | BK20.50m | 50 | | | | | 4.8 | 7.87 | 73.8 | 76.1 | 132.6 | 78.6 | 161996 | 105383 | 806 | 1168 | 229 | 0.91 | 0.53 |
| | | BK20.100m | 100 | | | | | 4.5 | 7.86 | 73.3 | 77.4 | 130.4 | 80.6 | 102618 | 89102 | 752 | 868 | 237 | 0.9 | 0.55 |
| | | BK20.200m | 200 | | | | | 4 | 7.74 | 72.9 | 74.8 | 131.8 | 77 | 101431 | 77741 | 727 | 727 | 229 | 0.83 | 0.48 |
| | | BK20.500m | 500 | | | | | 4.8 | 7.62 | 73.3 | 76.7 | 117.6 | 80.3 | 340083 | 275886 | 1261 | 1515 | 357 | 0.87 | 0.5 |
| | | BK20.850m | 850 | | | | | 6.5 | 7.71 | 75.9 | 74.1 | 88.5 | 68 | 92074 | 86915 | 621 | 674 | 366 | 0.87 | 0.56 |
| | BK14 | BK14.5m | 5 | 01.07.2017 | 54.09.926 N | 109.31.465 E | 129 | 5.5 | 7.63 | 78.1 | 68.3 | 118.9 | 84 | 174174 | 171428 | 663 | 274 | 268 | 0.93 | 0.67 |
| | BK13 | BK13.5m | 5 | 01.07.2017 | 53°53'53.10"N | 109°1'25.98"E | 13 | 6.2 | 7.9 | 84 | 66 | 80.9 | 65 | 94831 | 85391 | 851 | 1237 | 307 | 0.95 | 0.62 |
| | BK09 | BK09.5m | 5 | 30.06.2017 | 53°51'346" N | 108°42'911" E | 13 | 5.9 | 7.92 | 80.4 | 74.8 | 124.1 | 77 | 146124 | 126201 | 1079 | 1264 | 323 | 0.92 | 0.58 |
| | BK06 | BK06.5m | 5 | 30.06.2017 | 53°50'658" N | 108°40'195" E | 156 | 4.8 | 7.92 | 76.1 | 78 | 113.9 | 80 | 88294 | 78071 | 1217 | 991 | 378 | 0.9 | 0.53 |
| | | BK06.25m | 25 | | | | | 4.4 | 7.91 | 76.3 | 75.4 | 118 | 80 | 49540 | 46572 | 664 | 659 | 282 | 0.92 | 0.57 |
| | | BK06.50m | 50 | | | | | 5.1 | 7.82 | 78.9 | 778.7 | 115 | 82 | 233612 | 207530 | 1919 | 1477 | 414 | 0.94 | 0.6 |
| | | BK06.100m | 100 | | | | | 4.6 | 7.85 | 77.6 | 73.1 | 109.1 | 75 | 146124 | 102945 | 1000 | 1276 | 277 | 0.92 | 0.57 |
| | BK11 | BK11.5m | 5 | 30.06.2017 | 53°46'12.03"N | 109°5'9.56'E | 10 | 15.3 | 8.33 | 82.2 | 76.7 | 136.3 | 83.1 | 50396 | 35832 | 567 | 585 | 332 | 0.93 | 0.58 |
| Central | BK05 | BK05.5m | 5 | 29.06.2017 | 53°31'096" N | 108°24'583" E | 1512 | 5.5 | 7.9 | 77.1 | 78.7 | 134.3 | 80 | 115229 | 107228 | 811 | 1033 | 236 | 0.85 | 0.47 |
| | | BK05.25m | 25 | | | | | 4.1 | 7.81 | 74.1 | 77.4 | 134.8 | 77 | 48005 | 44526 | 543 | 478 | 228 | 0.73 | 0.41 |
| | | BK05.50m | 50 | | | | | 4.2 | 7.81 | 73.8 | 78 | 134.6 | 76.6 | 103557 | 88299 | 1067 | 1001 | 309 | 0.84 | 0.46 |
| | | BK05.100m | 100 | | | | | 4.2 | 7.81 | 74.3 | 78 | 134.7 | 77 | 96709 | 80346 | 1222 | 1240 | 389 | 0.89 | 0.5 |
| | | BK05.200m | 200 | | | | | 4 | 7.84 | 74.2 | 76.7 | 133.3 | 70.5 | 129517 | 74958 | 236 | 103 | 114 | 0.94 | 0.7 |
| | | BK05.500m | 500 | | | | | 3.7 | 7.7 | 74.4 | 75.4 | 127.3 | 71.5 | 160563 | 155621 | 456 | 305 | 151 | 0.82 | 0.51 |
| | | BK05.1000m | 1000 | | | | | 3.7 | 7.55 | 73.6 | 73.5 | 122.7 | 63.5 | 39227 | 36443 | 362 | 306 | 229 | 0.81 | 0.46 |
| | | BK05.1400m | 1400 | | | | | 3.6 | 7.7 | 74.5 | 80 | 114 | 77 | 66984 | 62868 | 373 | 250 | 185 | 0.82 | 0.49 |
| | BK22 | BK22.5m | 5 | 04.07.2017 | 53°23'524" N | 107°53'094" E | 592 | 4.9 | 7.86 | 74.7 | 76.5 | 86 | 83.8 | 170009 | 135771 | 1050 | 1159 | 281 | 0.82 | 0.49 |
| | | BK22.50m | 50 | | | | | 5.2 | 7.85 | 75.8 | 74.5 | 80.6 | 70.8 | 87940 | 85032 | 879 | 813 | 337 | 0.94 | 0.63 |
| | | BK22.200m | 200 | | | | | 7.8 | 7.77 | 75.2 | 72.2 | 72.8 | 68.3 | 143354 | 133777 | 825 | 701 | 264 | 0.93 | 0.62 |
| | | BK22.550m | 550 | | | | | 4.6 | 7.76 | 77 | 76.7 | 29.6 | 114 | 102066 | 97459 | 649 | 634 | 271 | 0.85 | 0.53 |
| | BK04 | BK04.5m | 5 | 29.06.2017 | 53°14'596" N | 108°24'583" E | NA | 9.5 | 8.47 | 87.3 | 72.8 | 44.4 | 80.6 | 62438 | 55549 | 726 | 838 | 311 | 0.89 | 0.53 |
| | BK24 | BK24.5m | 5 | 05.07.2017 | 53°00'525" N | 106°53'659" E | 35 | 7.8 | 8.11 | 81.3 | 76.7 | 31.8 | 98 | 131337 | 114477 | 743 | 685 | 248 | 0.85 | 0.48 |
| | BK03 | BK03.5m | 5 | 28.06.2017 | 52°41'401" N | 106°44'208" E | 1083 | 8.6 | 7.74 | 84 | 78.7 | 176.4 | 83 | 120296 | 107489 | 544 | 511 | 149 | 0.57 | 0.31 |
| | | BK03.25m | 25 | | | | | 4.8 | 7.67 | 75.5 | 77.4 | 179.9 | 82.6 | 132170 | 109849 | 915 | 879 | 279 | 0.87 | 0.5 |
| | | BK03.50m | 50 | | | | | 4.7 | 7.63 | 75.2 | 78.7 | 181.5 | 80.3 | 106227 | 97251 | 798 | 791 | 250 | 0.9 | 0.52 |
| | | BK03.100m | 100 | | | | | 4.3 | 7.6 | 74.7 | 76.1 | 183.2 | 77.6 | 105172 | 96013 | 1065 | 831 | 320 | 0.89 | 0.55 |
| | | BK03.200m | 200 | | | | | 4.3 | 7.6 | 74.6 | 74.8 | 184 | 73.3 | 59072 | 57656 | 344 | 107 | 172 | 0.91 | 0.63 |
| | | BK03.500m | 500 | | | | | 4 | 7.45 | 74.5 | 76.7 | 189.6 | 71.3 | 72297 | 69907 | 363 | 383 | 188 | 0.73 | 0.39 |
| | | BK03.1000m | 1000 | | | | | 4.8 | 7.48 | 79.1 | 79.3 | 195 | 68 | 144322 | 140470 | 436 | 301 | 156 | 0.7 | 0.38 |
| South | BK01 | BK01.5m | 5 | 28.06.2017 | 52°15'70" N | 106°02'90" E | 105 | 12.2 | 7.98 | 95.2 | 80.6 | 204.4 | 94.5 | 68506 | 59443 | 812 | 655 | 336 | 0.8 | 0.51 |
| | | BK01.25m | 25 | | | | | 8.1 | 7.56 | 86.8 | 83.2 | 224 | 83.5 | 124684 | 114488 | 1198 | 565 | 347 | 0.8 | 0.43 |
| | | BK01.50m | 50 | | | | | 8 | 7.54 | 78.9 | 74.5 | 234.3 | 26 | 89182 | 76475 | 1043 | 743 | 340 | 0.87 | 0.5 |
| | | BK01.85m | 85 | | | | | 6.5 | 7.88 | 83.5 | 83.2 | 205.6 | 87.7 | 59454 | 44623 | 685 | 640 | 306 | 0.91 | 0.57 |
| | BK02 | BK02.5m | 5 | 28.06.2017 | 52°15'70" N | 106°02'90" E | 10 | 13.7 | 7.62 | 97.5 | 80.6 | 202 | 83.3 | 77234 | 60742 | 647 | 533 | 296 | 0.84 | 0.48 |
| | BK26 | BK26.5m | 5 | 05.07.2017 | 51°52'638" N | 105°15'294" E | 1408 | 8.7 | 8.18 | 83.6 | 78.7 | 134 | 87.1 | 129657 | 103961 | 727 | 864 | 218 | 0.77 | 0.39 |
| | | BK26.25m | 25 | | | | | 6.5 | 8.06 | 76.1 | 76.7 | 138.2 | 86.1 | 142012 | 128621 | 567 | 514 | 157 | 0.47 | 0.28 |
| | | BK26.50m | 50 | | | | | 4.7 | 7.87 | 74.4 | 78 | 131 | 80.2 | 188707 | 150315 | 1204 | 954 | 312 | 0.88 | 0.53 |
| | | BK26.100m | 100 | | | | | 4.6 | 7.84 | 73.9 | 78 | 130.1 | 81.4 | 144537 | 114062 | 1102 | 977 | 343 | 0.91 | 0.54 |
| | | BK26.200m | 200 | | | | | 4.1 | 7.8 | 73.3 | 79.3 | 124.8 | 82.6 | 228022 | 188147 | 1301 | 828 | 326 | 0.91 | 0.56 |
| | | BK26.500m | 500 | | | | | 4 | 7.71 | 73.2 | 78.7 | 119.9 | 78.5 | 72429 | 55468 | 693 | 691 | 306 | 0.86 | 0.51 |
| | | BK26.1000m | 1000 | | | | | 3.9 | 7.64 | 73 | 78.7 | 109.5 | 82.8 | 36746 | 31456 | 324 | 299 | 204 | 0.87 | 0.5 |
| | | BK26.1350m | 1350 | | | | | 3.9 | 8.13 | 79 | 85.2 | 80.2 | 97.2 | 114918 | 109195 | 378 | 459 | 157 | 0.82 | 0.45 |
| | BK28 | BK28.5m | 5 | 06.07.2017 | 51°47'817" N | 104°46'449" E | 1381 | 11.7 | 8.28 | 89 | 76.7 | 90.7 | 88 | 197087 | 166833 | 830 | 1102 | 177 | 0.79 | 0.43 |
| | | BK28.25m | 25 | | | | | 5.6 | 8.14 | 77.1 | 71.5 | 95.3 | 104 | 321169 | 235066 | 2973 | 1993 | 698 | 0.94 | 0.59 |
| | | BK28.50m | 50 | | | | | 7 | 7.9 | 78 | 70 | 98.3 | 70 | 70786 | 53923 | 965 | 755 | 361 | 0.89 | 0.53 |
| | | BK28.100m | 100 | | | | | 4.6 | 7.87 | 75.2 | 77.4 | 94.2 | 85.4 | 111378 | 84474 | 1042 | 866 | 349 | 0.84 | 0.46 |
| | | BK28.200m | 200 | | | | | 4.6 | 7.8 | 75.4 | 77.3 | 107.8 | 83.7 | 81308 | 64784 | 993 | 1075 | 353 | 0.89 | 0.51 |
| | | BK28.500m | 500 | | | | | 4.9 | 7.69 | 76.1 | 78 | 96.9 | 85 | 133522 | 118083 | 943 | 1257 | 297 | 0.92 | 0.54 |
| | | BK28.1000m.CT | 1000 | | | | | 6.4 | 7.77 | 78.7 | 76.7 | 87.3 | 77 | 3234 | 3022 | 9 | 10 | 9 | 0.82 | 0.8 |
| | | BK28.1300m.CT | 1300 | | | | | 4.8 | 8.02 | 80.7 | 81.3 | 45.6 | 112 | 111545 | 107430 | 220 | 101 | 94 | 0.7 | 0.38 |

**Supplementary Table 2.** Identification, phylogenetic affinity and abundance of eukaryotic OTUs identified in Lake Baikal plankton.

(too large to be displayed in pdf; available in excel format)

Supplementary Table 3. Diversity, abundance and distribution of eukaryotic lineages previously thought to be exclusively marine identified in Lake Baikal plankton.

| Phylogenetic group | | Number of OTUs | Number of reads | No. samples where they occur |
|---|---|---|---|---|
| Bolidophyceae | | 85 | 9442 | 58 |
| Diplonema | | 4 | 505 | 21 |
| Radiolaria | | 2 | 22 | 1 |
| Syndiniales | | 73 | 1098 | 58 |
| All MAST | | 95 | 23722 | 61 |
| Specific MAST clades | MAST1 | 1 | 2 | 1 |
| | MAST2 | 22 | 8250 | 54 |
| | MAST3 | 13 | 1296 | 41 |
| | MAST4 | 6 | 219 | 10 |
| | MAST6 | 4 | 3453 | 50 |
| | MAST8 | 1 | 4 | 3 |
| | MAST12 | 47 | 10487 | 59 |
| | MAST20 | 1 | 11 | 1 |

Supplementary Table 4. ANOSIM analyses between pairs of Lake Baikal plankton sample groups defined as a function of depth, basin of origin and coastal vs. pelagic location. ANOSIM were calculated upon 999 permutations between pairs of sample groups.

| | R | p-value |
|---|---|---|
| Epi-Meso | 0.3821 | 0.0001 |
| Epi-Bathy | 0.7262 | 0.0001 |
| Meso-Bathy | 0.7262 | 0.0001 |
| Southern-Central | 0.09237 | 0.0001 |
| Southern-North | 0.2201 | 0.0001 |
| Central-North | 0.1442 | 0.0001 |
| Coastal-Pelagic | 0.05955 | 0.2339 |

Supplementary Table 5. Tentative classification of phylogenetic lineages within broad functional categories.

| Phylum | Super_Group | Lifestyle |
|---|---|---|
| Chlorophyta | Archaeplastida | Autotrophs |
| Cryptophyta | Hacrobia | Autotrophs |
| Dinophyta | Alveolata | Autotrophs |
| Haptophyta | Hacrobia | Autotrophs |
| Ochrophyta | Stramenopiles | Autotrophs |
| Rhodophyta | Archaeplastida | Autotrophs |
| Charophyta | Archaeplastida | Autotrophs |
| Apusozoa | Apusozoa | Free-living heterotrophs |
| Bicoecea | Stramenopiles | Free-living heterotrophs |
| Breviatea | Amoebozoa | Free-living heterotrophs |
| Centroheliozoa | Hacrobia | Free-living heterotrophs |
| Cercozoa | Rhizaria | Free-living heterotrophs |
| Choanoflagellida | Opisthokonta | Free-living heterotrophs |
| Ciliophora | Alveolata | Free-living heterotrophs |
| Conosa | Amoebozoa | Free-living heterotrophs |
| Discoba | Excavata | Free-living heterotrophs |
| Fungi | Opisthokonta | Free-living heterotrophs |
| Katablepharidophyta | Hacrobia | Free-living heterotrophs |
| Labyrinthulea | Stramenopiles | Free-living heterotrophs |
| Lobosa | Amoebozoa | Free-living heterotrophs |
| MAST | Stramenopiles | Free-living heterotrophs |
| Metamonada | Excavata | Free-living heterotrophs |
| Radiolaria | Rhizaria | Free-living heterotrophs |
| Telonemia | Hacrobia | Free-living heterotrophs |
| Picozoa | Uncertain | Free-living heterotrophs |
| Mesomycetozoa | Opisthokonta | Free-living heterotrophs |
| Pirsonia | Stramenopiles | Free-living heterotrophs |
| Syndiniales | Alveolata | Putative parasites |
| Cryptomycota | Opisthokonta | Putative parasites |
| Hyphochytriomyceta | Stramenopiles | Putative parasites |
| Oomycota | Stramenopiles | Putative parasites |
| Apicomplexa | Alveolata | Putative parasites |
| Opalinata | Stramenopiles | Putative parasites |
| Perkinsea | Alveolata | Putative parasites |
| Aphelida | Opisthokonta | Putative parasites |
| Uncertain_Alveolata | Alveolata | Uncertain |
| Uncertain_Amoebozoa | Amoebozoa | Uncertain |
| Uncertain_Opisthokonta | Opisthokonta | Uncertain |
| Uncertain_Stramenopiles | Stramenopiles | Uncertain |

Supplementary Table 6. Functional and phylogenetic classification of OTUs used for network construction. Only OTUs that were present in more than 20% of samples with a relative abundance higher than 0.01% were considered.

| | Epipelagic | | Mesopelagic | | Bathypelagic | |
|---|---|---|---|---|---|---|
| | Not connected | Connected | Not connected | Connected | Not connected | Connected |
| Functional classification | | | | | | |
| Autotrophs | 69 | 45 | 104 | 29 | 59 | 23 |
| Heterotrophs | 35 | 51 | 59 | 25 | 74 | 9 |
| Parasites | 5 | 1 | 9 | 1 | 7 | 1 |
| Phylogenetic ascription | | | | | | |
| Apicomplexa | 1 | 0 | 1 | 0 | 1 | 0 |
| Apusozoa | 0 | 0 | 0 | 0 | 0 | 0 |
| Bicoecea | 2 | 1 | 1 | 0 | 3 | 0 |
| Centroheliozoa | 0 | 0 | 0 | 0 | 0 | 0 |
| Cercozoa | 13 | 9 | 12 | 8 | 13 | 3 |
| Chlorophyta | 9 | 2 | 13 | 2 | 7 | 1 |
| Choanoflagellida | 0 | 2 | 2 | 2 | 4 | 0 |
| Ciliophora | 11 | 33 | 23 | 8 | 33 | 2 |
| Conosa | 0 | 0 | 1 | 0 | 1 | 0 |
| Cryptomycota | 3 | 0 | 2 | 0 | 3 | 1 |
| Cryptophyta | 11 | 3 | 8 | 3 | 13 | 0 |
| Dinophyta | 13 | 6 | 15 | 6 | 8 | 3 |
| Discoba | 0 | 0 | 0 | 1 | 1 | 0 |
| Fungi (+Aphelida) | 5 | 1 | 10 | 3 | 11 | 3 |
| Haptophyta | 5 | 2 | 3 | 1 | 3 | 1 |
| Hyphochytriomyceta | 0 | 0 | 0 | 0 | 0 | 0 |
| Katablepharidophyta | 1 | 0 | 0 | 1 | 1 | 0 |
| Labyrinthulea | 0 | 0 | 0 | 0 | 0 | 0 |
| Lobosa | 0 | 0 | 0 | 0 | 0 | 0 |
| MAST | 3 | 2 | 8 | 1 | 4 | 1 |
| Mesomycetozoa | 0 | 0 | 1 | 0 | 1 | 0 |
| Ochrophyta | 31 | 32 | 65 | 17 | 28 | 18 |
| Oomycota | 1 | 0 | 3 | 0 | 2 | 0 |
| Opalinata | 0 | 0 | 0 | 0 | 0 | 0 |
| Perkinsea | 0 | 1 | 3 | 0 | 1 | 0 |
| Picozoa | 0 | 0 | 0 | 0 | 0 | 0 |
| Pirsonia | 0 | 1 | 0 | 0 | 0 | 0 |
| Radiolaria | 0 | 0 | 0 | 0 | 0 | 0 |
| Syndiniales | 0 | 0 | 0 | 0 | 0 | 0 |
| Telonemia | 0 | 2 | 1 | 2 | 2 | 0 |
| Uncertain_Alveolata | 0 | 0 | 0 | 0 | 0 | 0 |
| Uncertain_Opisthokonta | 0 | 0 | 0 | 0 | 0 | 0 |
| Uncertain_Stramenopiles | 0 | 0 | 0 | 0 | 0 | 0 |

Supplementary Table 7. Properties of networks built upon Lake Baikal plankton OTUs for the three depth categories of the water column. Only OTUs that were present in more than 20% of samples with a relative abundance higher than 0.01% were considered.

| | Eepipelagic | | Mesopelagic | | Bathypelagic | |
|---|---|---|---|---|---|---|
| Parameter | Basin | % | Basin | % | Basin | % |
| Positive edges | 10 | 4.8 | 1 | 1.61 | 1 | 1.26 |
| Negative edges | 198 | 95.19 | 61 | 98.38 | 78 | 98.73 |
| Total number of edges | 208 | | 62 | | 79 | |
| Total number of nodes | 206 | | 227 | | 173 | |
| Connected nodes | 97 | 47.08 | 55 | 24.22 | 33 | 19.07 |
| Connectance | 0.04 | | 0.041 | | 0.14 | |
| Clustering coefficient | 0.31 | | 0.25 | | 0.43 | |
| Most connected node | X406443 (27) | | X403371 (9) | | X401791, X403173 (18) | |
| Average path length | 3.51 | | 2.5 | | 2.11 | |
| Mean node degree | 2.02 | | 0.55 | | 0.91 | |
